# Characterisation of the historic demographic decline of the British European polecat population

**DOI:** 10.1101/2024.10.30.621102

**Authors:** R. Shaw, J. MacPherson, A. C. Kitchener, G. J. Etherington, W. Haerty

**Affiliations:** Earlham Institute, Norwich Research Park, Colney Lane, Norwich, NR4 7UZ; University of East Anglia, Norwich Research Park, Norwich, NR4 7TJ; Vincent Wildlife Trust, Ledbury, Herefordshire, HR8 1EP; 4 Department of Natural Sciences, National Museums Scotland, Edinburgh EH1 1JF, UKUK and School of Geosciences, University of Edinburgh, Drummond Street, Edinburgh EH8 9XP, UK

**Keywords:** population genomics, demographic history, range expansion

## Abstract

The European polecat (*Mustela putorius*) has a widespread distribution across many countries of mainland Europe but is documented to be declining within these ranges. In Britain, direct persecution led to a severe decline of the polecat population during the 19^th^ century. Unlike European mainland populations, it is now recovering across much of its former British range. The genomic and conservation implications of such a severe demographic decline, followed by the current recovery, have still to be characterised in the European polecat in Britain. Here we carry out population-level whole-genome analyses of 65 polecats from Britain (Wales and England) and the European mainland. Our analyses reveal that Welsh polecats show genetic variability from both English and European polecats, while British polecats as a whole exhibit signs of genetic isolation from mainland European populations. We also reconstructed the demographic history of the Welsh polecat to quantify the magnitude of the bottleneck. Our analyses confirmed the drastic decline of the Welsh polecat’s effective population size, with a severe genetic bottleneck around 30-40 generations ago (1854-1894). We investigated whether whole genome diversity reflected this demographic event and found that Welsh polecats had significantly less genetic diversity than English polecats, but not European polecats. Runs of homozygosity and genetic load present in Welsh and English polecat genomes also indicated recent historic inbreeding. Our findings suggest that the increase in the British polecat population size may be attributed to admixture events. Additionally, we demonstrate that the Welsh polecat constitutes a genetically distinct population, which could be crucial for the overall conservation of European polecats by preserving unique genetic diversity.

## 1.0 Introduction

Human-mediated actions are considered some of the most serious threats to the survival of small carnivore populations (Willcox 2020; C. Marneweck et al. 2021; C. J. Marneweck et al. 2022; Prugh et al. 2023). This includes active persecution and habitat fragmentation, which reduces connectivity between individuals, thereby limiting gene flow and further contributing to reducing effective population size (N_e_) (Croose et al. 2018; C. Marneweck et al. 2021). The ensuing decline in genetic diversity further decreases future resilience to arising challenges including environmental changes (Frankham et al. 2017). Smaller populations are therefore highly susceptible to inbreeding and genetic drift, which can limit the species’ ability to purge deleterious mutations due to reduced efficacy of purifying selection at low N_e_, coupled with increasing efficacy of drift which increases their probability of fixation. This phenomenon allows for weakly deleterious mutations to segregate at higher frequencies which can negatively affect the population and lead to an increased risk of extinction (Lande 1994; Spielman, Brook, and Frankham 2004; Khan et al. 2021).

Similarly, populations that have recently undergone rapid expansion often display genetic signatures of low effective population size, partly due to allele surfing, where certain gene variants increase in frequency among individuals at the forefront of the expansion (Welles and Dlugosch 2018; Cisternas-Fuentes and Koski 2023; Excoffier and Ray 2008; Graciá et al. 2013). Likewise, the ‘founder effect’ can leave a distinct genetic imprint by reducing genetic diversity, often resulting in greater genetic differentiation between populations (Szewczyk et al. 2019; Jarausch et al. 2021). These signals are often associated with historical (post-glacial), contemporary (often human-mediated) introductions of species (Sendell-Price, Ruegg, and Clegg 2020) or by range expansions. The latter has been well documented in the recent, rapid range expansion of the grey wolf (*Canis lupus*) across Europe, where recolonisation of these areas by expanding populations can lead to altering genetic structure across the landscape (Gustafson et al. 2018; Szewczyk et al. 2019; Jarausch et al. 2021).

Habitat fragmentation or population displacement can also threaten the genetic integrity of species through introgression and hybridisation. As the boundaries between species populations overlap, or through the introduction of invasive species, populations may become more susceptible to interbreeding with closely related sister species. Hybridisation with closely related taxa has been shown to have negative consequences, particularly in cases for endangered species (Allendorf et al. 2001). These arise through outbreeding depression which causes a reduction in the fitness of offspring (P. W. Hedrick et al. 2019) or genomic swamping whereby parental genetics are overwhelmed by the hybrid, potentially leading to their extinction (Howard-McCombe et al. 2023). Hybridisation can also impact how a species is managed, with there often being uncertainty over the classification and protection of hybrids in the wild (Senn et al. 2019; Moroni et al. 2022).

Adaptive introgression, which occurs through successful hybridisation between two species, can be particularly beneficial for populations recovering from a bottleneck (Philip W. Hedrick 2013). In such cases, genetic diversity is low, and introgression can introduce new variants into the offspring’s genome. This phenomenon has been observed in the Alpine ibex (*Capra ibex*) (Brambilla et al. 2024) and Scottish wildcat (*Felis silvestris*) (Howard-McCombe et al. 2023), where adaptive introgression has replenished genetic diversity in previously depleted genomic regions, such as immune-related genes, thereby increasing the populations’ adaptive potential. This can be beneficial for recovering populations, yet adds to the complexity of managing hybrid individuals, as distinguishing signals of introgression are complex and require well-defined local ancestry estimates (Howard-McCombe et al. 2023; Brambilla et al. 2024).

Therefore, it is crucial to monitor and investigate genomic signals to understand how populations adapt to and recover from environmental change. This knowledge can inform conservation strategies, such as prioritising populations for protection (von Takach et al. 2024), monitoring reintroductions (Gajdárová et al. 2023) or translocations (Farquharson et al. 2022), identifying hybrid individuals to inform action plans (Senn et al. 2019) and contributing to ex-situ breeding programmes (Speak et al. 2024). Additionally, capturing the temporal fluctuations of genomic changes in a recovering population undergoing range expansion is essential to inform conservation management strategies to prevent long-term genetic fragmentation and monitor loss (or gain) of adaptive potential (Thomas et al. 2022).

Several of the aforementioned environmental threats are known to severely impact populations of the European polecat (*Mustela putorius*), a medium-sized carnivore of the Mustelidae family. This family is a great example of adaptive radiation, as species belonging to this family occupy a wide range of niches within different climates and habitats (Schluter 2000; Koepfli et al. 2008). Despite this, and despite the polecat inhabiting a wide geographical range across mainland Europe (Virgós 2003; Mestre, Ferreira, and Mira 2007; Brzeziński, Marzec, and Żmihorski 2010), in several of these countries their populations are listed as or suspected to be declining (Maran et al. 2016; Croose et al. 2018) due to habitat fragmentation, prey availability, poisoning, and persecution in continental Europe (Eeraerts et al. 2022; Barrientos et al. 2024; Szapu et al. 2024). Hybridisation has also been known to occur between the European polecat and other closely related mustelid species, such as the Steppe polecat (*Mustela eversmanii*) and the severely endangered European mink (*Mustela lutreola*) (Cabria et al. 2011; Cserkész et al. 2021).

### Polecat population in Britain

In Britain, the European polecat has undergone an extensive population reduction and range contraction, followed by a subsequent rapid range and population expansion over the last 70 years (Langley and Yalden 1977; Birks and Kitchener 1999; S. 2008; Costa et al. 2013; Croose 2016). The population experienced a severe decline in numbers during the late 19th century as the species was persecuted heavily by gamekeepers and regarded as pests which impacted gamebird populations (Birks and Kitchener 1999; Costa et al. 2013). This led to a contraction in the species range, with small pockets of populations in Cumbria and Scotland persisting until the early 20th century and a stronghold in central Wales and the neighbouring English border counties of Herefordshire and Shropshire (Langley and Yalden 1977; Harris et al. 1995).

The existing stronghold in Wales started to recover in population size after the end of the First World War This resulted in a period of reduced persecution and coincided with the recovery of rabbit populations in the post myxomatosis period, a key prey species in the polecats diet (Langley and Yalden 1977). To aid its protection, the polecat was listed on schedule 6 of the UK Wildlife and Countryside Act in 1981, and since 2007 has been listed as a priority species on the UK Biodiversity Action Plan (BRIG 2007). As the population began to expand eastwards across its former range, it came into contact with populations of feral domestic ferrets (*Mustela putorius furo*) (Davison et al. 1999; Costa et al. 2013; Etherington et al. 2022). Ferrets are thought to be derived from the European polecat through domestication that occurred over 2000 years ago (Blandford 1987; Davison et al. 1999) and have long been used for hunting purposes. In Britain, they are known to have been used for predominantly hunting rabbits, and are thought to have been released into the wild and accidentally introduced. Owing to the short evolutionary distance between the two species, the European polecat and the domestic ferret readily hybridise and this has been well-documented to occur in Britain (Costa et al. 2013; Etherington et al. 2022).

Previous work on the British population of polecats has focussed on assessing genome introgression between European polecats and feral domestic ferrets (Davison et al. 1999; Costa et al. 2013; Etherington et al. 2022) or has aimed to model their demographic history (Costa et al. 2013). These studies identified high degrees of genome introgression in British polecats outside of their stronghold, emphasising the importance of the Welsh population for conservation of the British European polecat. However, genetic assessments for the Welsh population have either been conducted using microsatellites and mitochondrial genes, which did not provide enough power to confidently assess the size of the genetic bottleneck (Costa et al. 2013), or were limited in the number of Welsh individuals used in their assessment (Etherington et al. 2022).

Here, we are able to characterise the demographic history of the British European polecat population through using whole genome resequencing of English, Welsh and mainland European polecats. We also further mine the genome for signatures of the population bottleneck, documenting what impact the decline and rapid expansion had on the population and assessing the conservation implications for British polecats in comparison to mainland European polecat populations.

## 2.0 Methods

### 2.1 Sample Collection

For this study, 31 roadkill British polecat samples were resourced from the Vincent Wildlife Trust 2014-2015 survey (Croose 2016) (Figure 1b). Carcasses were collected by members of the public who were requested to freeze the carcass where possible. These were then sent to the Centre for Ecology and Hydrology (CEH) via a pre-paid postage box and the carcasses were frozen upon receipt. Samples were selected based on geographical location from areas around Wales and from a trajectory across England from west to east to capture any diversity across the predicted range expansion of the polecat in Britain (Figure 1c), (Supplementary table sheet 1 (S1)). The home ranges of polecats in Britain are known to vary seasonally, and male polecats tend to disperse further than females (Birks and Kitchener 1999), with males travelling as far as 8 km between dens (Blandford 1987). To avoid incorporating familial bias into our selection through potentially selecting related individuals, we selected carcasses that were located greater than 10 km apart. We further increased our sample size through the inclusion of publicly available data consisting of eight domestic ferrets genomes, 19 British polecats, and 15 European mainland polecat samples (Etherington et al. 2022).

**Figure 1.**
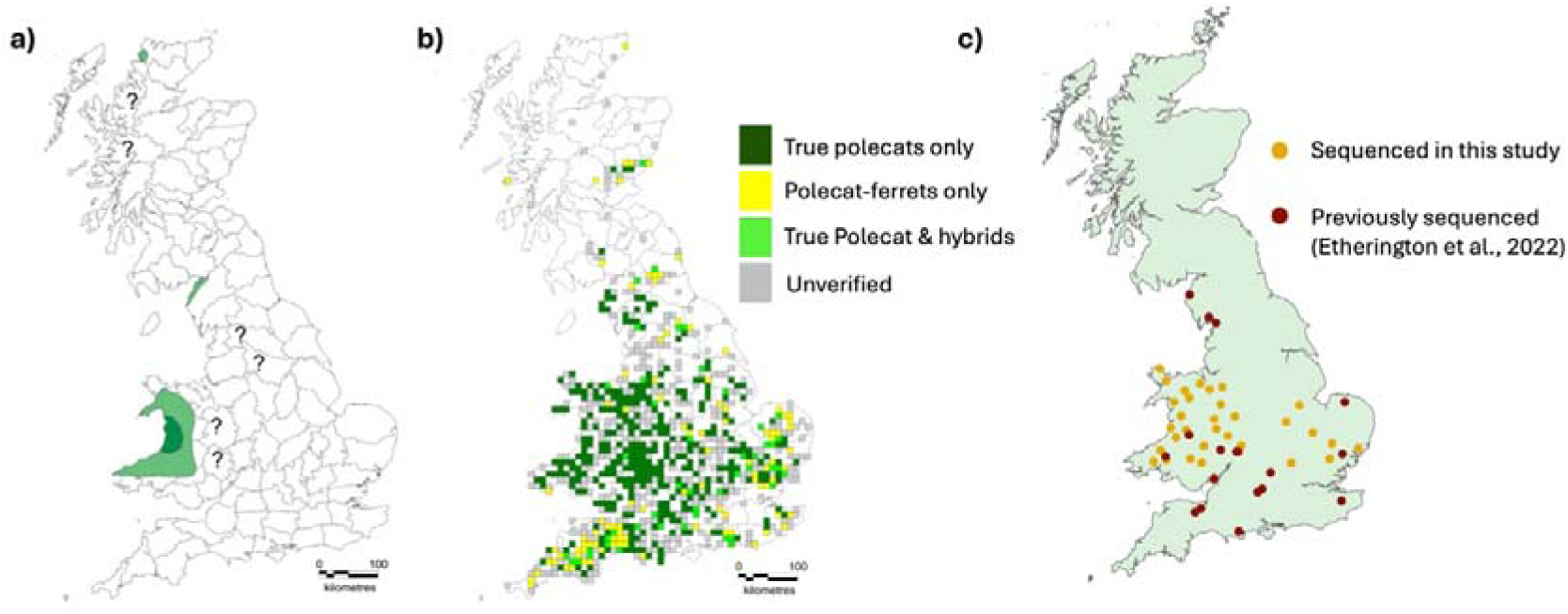
(a) The range of the European polecat in Britain in 1915. Dark green areas indicate the population stronghold, light green indicates known occurrences, and “?” refers to uncertain occurrences of polecats in those areas (Langley and Yalden 1977). (b) Records received during the Vincent Wildlife Trust 2014-15 survey of verifiable sightings, based on their phenotype, of true polecats (dark-green), polecat-ferret hybrids (yellow), both true polecats and hybrids (lime green), and unverifiable (grey) (Croose 2016). (c) Samples selected to be sequenced in this study (dark yellow) and (red) samples that were previously sequenced by (Etherington et al. 2022). Figures 1(a) and 1(b) were adapted from Croose, 2016.

### 2.2 DNA Extraction

DNA extraction from tissue samples was carried out using a Qiagen^TM^ DNEasy Blood & Tissue kit according to the manufacturer’s protocol. Samples were quality checked using a Nanodrop Spectrophore and DNA quantity was checked using a Qubit Fluorometer. Samples with a low yield, and with 260/280 and 260/230, absorbance values outside the desired range (260/280 <1.80, 260/230 <2.0) were further cleaned up using Beckman Coulter^TM^ AMPureXP Beads. A total of 31 samples were selected for library preparation and sent for sequencing. Figure 1c shows the distribution of the samples sent for sequencing and the location of British polecats already sequenced.

### 2.3 Library preparation & sequencing

Samples were sequenced by The Earlham Institute Technical Genomics with LITE library preparation (Perez-Sepulveda et al. 2021) and sequenced using an Illumina NovaSeq 6000 generating paired-end (2 x 150bp) reads. Samples were run on one lane with a predicted coverage of 10x.

### 2.4 Sequencing QC, Read mapping & SNP calling

Raw read quality for all British and European samples were first assessed using fastQC version 0.11.9 (Andrews 2010), and any reads below the threshold (Q20) were trimmed using TRIMMOMATIC version 0.39 (Bolger, Lohse, and Usadel 2014). Raw reads were mapped to a new version of the domestic ferret genome MusPutFur1.0 scaffolded with Bionano data (<u>GCA_920103865.1</u>) (Peng et al. 2014; Etherington et al. 2022) using BWA MEM version 0.7.17 (Li and Durbin 2009). Duplicate reads were then marked using PicardTools version 2.1.1 (“Picard Toolkit” 2019). Single nucleotide polymorphisms (SNPs) were then called using GATK version 4.5.0 (McKenna et al. 2010) following the best practice workflow for SNP discovery (van der Auwera and O’Connor 2020). Briefly, BaseRecalibrator was run on each sample bam file to detect and correct patterns of systematic errors in the base quality scores. The recalibration metric was then applied to each sample bam file, and HaplotypeCaller was run for each sample before consolidating all of them together using GATK GenomicsDBImport. For downstream analysis, BCFtools version 1.15.1 (Danecek et al. 2021) was used to select only biallelic SNPs with a quality score (QUAL) of at least 20 and a minor allele count of at least three (AC3). A further step of pruning variants over a linkage disequilibrium (LD) threshold of r2 >0.6 within 50kb windows was carried out using the bcftools prune command. The resulting SNP dataset was used for all downstream analyses unless stated. For GONE analyses, all variants called before pruning were used.

### 2.5 Relatedness

We assessed relatedness between samples using the KING algorithm in PLINK version 1.9 (Purcell et al. 2007; Chang et al. 2015). KING kinship coefficients determine first-degree relations (parent-child, full siblings) as values between 0.17-0.35, second-degree relations correspond to 0.08-0.17 and third-degree relations as between 0.04-0.08 (Manichaikul et al. 2010). One pairing between samples S17 and S0012_VWT665 had a kinship coefficient of 0.07, suggesting a third-degree level of relatedness. Both of these individuals were English polecats but were found in very different geographical locations (∼291km apart). We removed S17 from all of the downstream analysis as this sample was found in Waterrow, England, and was therefore outside of the west-to-east trajectory that we were hypothesising was the route of the Welsh polecat expansion.

### 2.6 Population Structure

We first generated a principal component analysis (PCA) using the whole-genome SNP dataset of 65 polecats and 8 domestic ferrets in PLINK version 1.9 (Purcell et al. 2007; Chang et al. 2015) to determine the British population structure. A Spearman correlation test was implemented in R version 4.1.1 (R Core Team 2024), using values from PC1 and sample longitude coordinates, to ascertain whether population structure in Britain could be correlated with geographical location. We used ADMIXTURE version 1.3 (Alexander, Novembre, and Lange 2009) to perform ancestry runs on the SNP dataset for *K* values 1-12, and a CV error was used to compare all *K* values. Replicates for each ancestry run were carried out using the custom python wrapper (admixturewrapper.py). Pairwise F_st_ values between populations were computed using VCFtools version 0.1.16 (Danecek et al. 2011). To further detect structure within the British population, we tested for isolation by distance between individuals. We did this by first calculating the genetic distance (Hamming distance) between British samples using PLINK version 1.9 (Purcell et al. 2007; Chang et al. 2015). We then used a geographic distance matrix between British samples using the geosphere package in R version 4.1.1 (R Core Team 2024; Hijmans 2024), inputting longitudinal and latitudinal coordinates to output distance in km. A mantel test was used to test for isolation by distance between the two matrices.

We used RAXML-ng version 0.9.0 (Stamatakis 2014) to infer a phylogenetic tree from our whole-genome SNP dataset. The phylogeny was generated using the GTRGAMMA model of evolution with the Lewis bias ascertainment correction and 100 rapid bootstraps.

### 2.7 Genomic Diversity

To assess the genetic diversity between populations, we used VCFtools version 0.1.16 (Danecek et al. 2011) to detect nucleotide diversity in 20 Mb windows across all samples. We tested for significant differences in the nucleotide diversity estimates between populations by using an analysis of variance test (ANOVA) implemented in R version 4.1.1(R Core Team 2024). Genome-wide SNPs that were private to one population, or that were shared across populations, were identified using VCFtools. The observed proportion of heterozygous sites and the inbreeding coefficient (f) for each individual per population were also calculated using VCFtools.

### 2.8 Detection of historic bottleneck: Runs of homozygosity and identification of deleterious mutations

To detect signatures associated with a genetic bottleneck, we calculated the distribution of runs of homozygosity (ROH) using PLINK. Since calling ROH can be sensitive to low sequencing depth in areas of the genome, which can introduce genotype errors (Ceballos et al. 2018), we only calculated ROH for individuals per population that had an average read depth of 6.0 or higher. We used the following recommended parameters to call ROH for low coverage samples - plink --geno 0.5 --maf 0.1 --homozyg --homozyg-density 50 --homozyg-gap 1000 --homozyg-kb 300 --homozyg-snp 50 --homozyg-window-het 4 --homozyg-window-missing 5 --homozyg-window-snp 50 --homozyg-window-threshold 0.05 --aec -- double-id (Ceballos et al. 2018). We then categorised ROH into three length classes; 300- 500kb, 500 kb-1Mb, and >1Mb.

SnpEff version 5.1 (Cingolani et al. 2012) was used to detect the functional impacts of the biallelic SNPs on the genome of the European polecat. SnpEff compares a set of variants to a known annotation database and compares the derived alleles within genic regions to predict the impact of that variant. In this case, we used the domestic ferret annotation as the input database. We assessed whether mutational load differed between populations and this was calculated based on homozygous and heterozygous mutational load within missense and loss-of-function (LoF) categories.

Due to the potential bias that could arise by aligning reads to the domestic ferret reference genome to infer deleterious mutations within the polecat genome landscape, we polarised our variant calls using the Steppe polecat (*Mustela eversmanii*) and weasel (*Mustela nivalis*) as outgroups to infer the ancestral allele at sites called by SnpEff version 5.1 (Cingolani et al. 2012) as deleterious.

To infer the ancestral allele within outgroups, we selected species from the Mustelidae family with available genomes: European polecat (*Mustela putorius*) (GCA_902207235.1) (Etherington et al. 2020), domestic ferret (*Mustela putorius furo*) (GCA_920103865.1) (Peng et al. 2014; Etherington et al. 2022), Steppe polecat (*Mustela eversmanii*) (GCA_963422785.1), Black-footed ferret (*Mustela nigripes*) (GCF_022355385.1) (Kliver et al. 2023), stoat (*Mustela erminea*) (GCA_009829155.1), stone marten (*Martes foina*) (GCA_040938555.1) (Tomarovsky et al. 2025), and Eurasian otter (*Lutra lutra*) (GCA_902655055) (Mead et al. 2020). Additionally, we included three mammalian outgroups: human (*Homo sapiens*) (GCA_000001405.15) (Schneider et al. 2017), domestic cat (*Felis catus*) (GCF_000181335.3) (Buckley et al. 2020), and domestic dog (*Canis lupus familiaris*) (GCF_011100685.1) (Wang et al. 2021) to construct a whole-genome alignment.

We generated a whole-genome alignment using Progressive Cactus v2.2.0 (Armstrong et al. 2020) without specifying branch lengths. Next, we extracted ancestral repeats from the alignment and inferred a phylogenetic tree under a neutral model using phyloFit (part of the PHAST toolkit) (Siepel and Haussler 2004). This tree was then used as a guide for rerunning Progressive Cactus to refine the alignment.

We subsequently extracted variants from this alignment for four sister species (domestic ferret, European polecat, Steppe polecat, and weasel), using the domestic ferret as the reference, via halSNPs v2.2 (Hickey et al. 2013). Finally, we intersected the extracted variants with our called SnpEff variants to identify alleles derived in the European polecat.

### 2.9 Historic effective population size

Recent historic effective population size (N_e_) was estimated using GONE (Santiago et al. 2020). GONE estimates recent effective population size by calculating LD between pairs of SNPs and finds the N_e_ that best fits the observed LD spectrum. GONE assumes that the population being tested is closed, with no recent admixture to allow for unbiased estimates in the calculation of effective population size. Therefore, we ran GONE for Welsh, English, and European populations separately. We restricted GONE runs for the English population to those individuals in the south, central and eastern regions of England. This was to account for any potential gene flow between individuals located within the border counties of Wales and Welsh individuals, as well as lack of gene flow due to the geographic isolation of individuals within the north of England. We further removed an individual (S0024_VWT1661) from the Welsh population run due to apparent recent hybridisation (F1 hybrid) identified by PCA and ADMIXTURE analyses. We accounted for low sequencing depth by further removing individuals from the analyses that had a mean sequencing depth at sites of less than 5x coverage. For each population, we simulated 2000 generations and calculated 400 bins. GONE takes up to 200 chromosomes as input data, so we removed scaffolds that overlapped with the X chromosome, using the annotated X chromosome of the black-footed ferret (*Mustela nigripes*) for reference (Kliver et al. 2023) and used the largest 100 autosomal scaffolds as input, constituting 78% of the whole genome assembly. GONE calculated LD for up to 10,000 SNPs per scaffold. Each simulation was run 100 times, with a new subsample of 10,000 SNPs for each run. For each population, we plotted the mean Ne for the 100 runs.

## 3.0 Results

### 3.1 Whole Genome Resequencing

We generated whole-genome sequence data for 31 European polecats from Wales and England. Samples had been chosen from the location of the known refugium (Figure 1a), across a trajectory of the historic range expansion following the population’s recovery from a genetic bottleneck. These were combined with an already published dataset of 34 European polecats from Italy (5), Germany (2), Austria (3), France (3), Spain (2), Wales (3) and England (16), along with 8 domestic ferrets. Sequence reads were aligned to the domestic Ferret genome. Initial exploratory analyses of called variants determined that there was a presence of batch effect (Lou and Therkildsen 2022) across the different sequencing runs, and steps taken to alleviate this effect are described in the Supplemetary Methods. Higher-coverage samples were downsampled to account for coverage biases, and SNPs were then called after downsampling (Supplementary table S2). We identified a total of 2,941,545 SNPs.

### 3.2 Population Structure Using Genomic Data

Principal component analysis (PCA) of whole-genome SNPs revealed 3 distinct clusters of samples of polecats from (1) mainland Europe, (2) Britain, and (3) domestic ferrets (Figure 2a). Within this, PCs 1 (26%) & 2 (15.8%) revealed the English and Welsh samples clustered together and away from mainland European samples. To assess structure within the British population, we repeated the PCA only using the 49 British samples, which revealed a geographical separation of samples from West (Welsh) and East (English) (Figure 2b). Longitudinal coordinates strongly correlated with PC1 (r_s_ = -0.754, p <0.01). Outliers to this relationship that clustered with the Eastern individuals are from northwestern and southern England, suggesting that Welsh samples were distinct from English samples.

**Figure 2.**
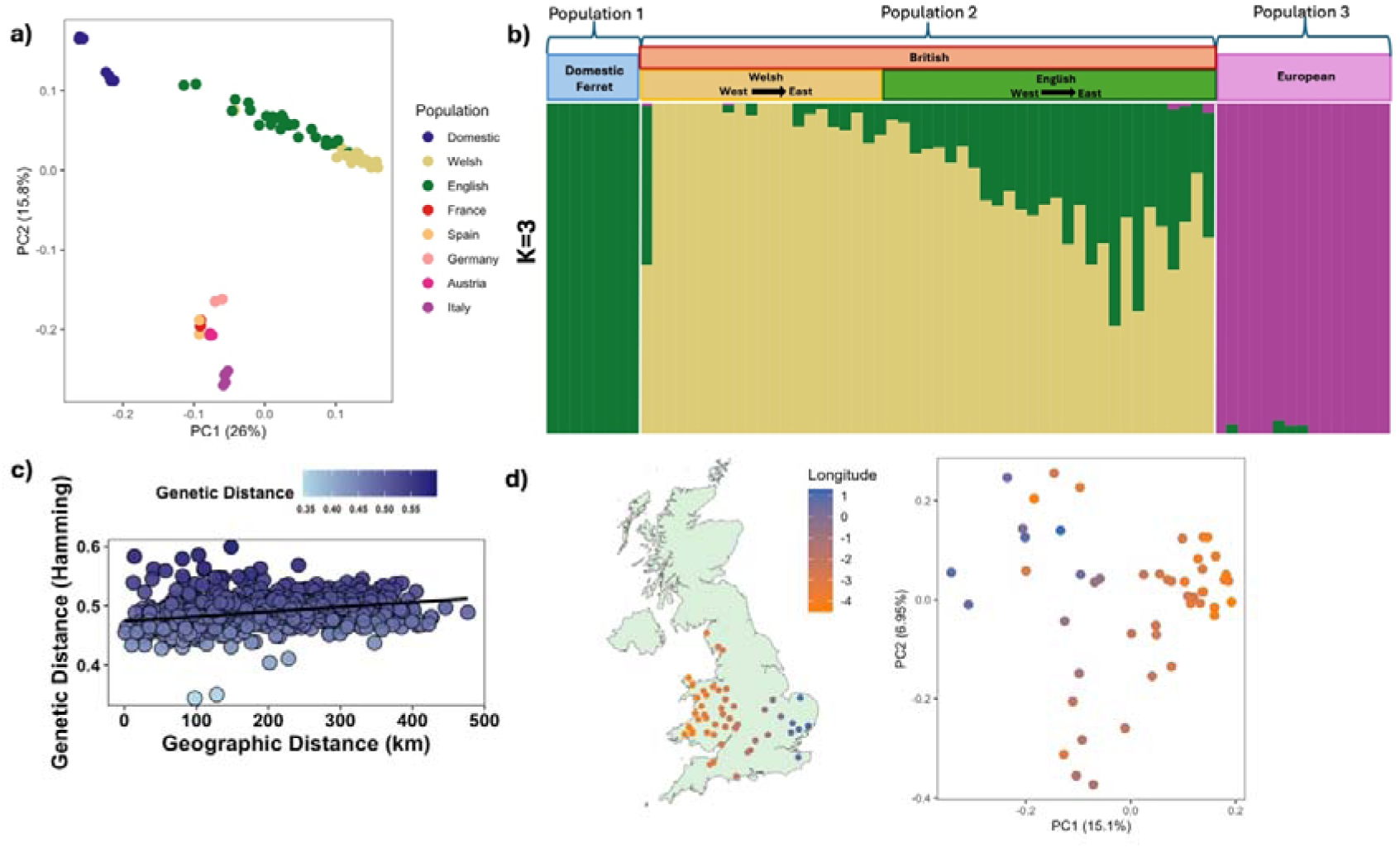
Population structure of the European polecat in Great Britain and across mainland Europe. (a) PCA based on whole genome SNPs for all European polecats and domestic ferrets. (b) Admixture results for whole-genome SNPs between the four groups (Domestic ferret, English, Welsh and mainland European polecats). (c) Isolation by distance of samples plotted using a pairwise Hamming genetic distance and geographic distance matrix (d) Map of Great Britain demonstrating longitude coordinates of samples and PCA based on whole genome SNPs for all British polecats, which is supported by a significant correlation between PC1 and longitude coordinates.

ADMIXTURE analyses were best supported by the cross-validation error of *K=3* (Supplementary Figure S3 & S5), indicating the existence of three genetically distinct populations across all of the sampled populations; domestic ferret, British (encompassing English and Welsh populations) and European polecats. These results appear to resolve the British population as a homogenous group distinct from the European mainland population, however this could also be due to the small sample size representing different European populations (Figure 2c). Under these parameters, the Welsh population demonstrated a lower degree of admixture with the domestic ferret than English individuals (Figure 2c) (Supplementary table S3), with the exception of one individual, which is likely a polecat-ferret hybrid (S0024_VWT1661). These results suggest that there is evidence of hybridisation and introgression occurring between English polecats and feral domestic ferrets at varying degrees across the polecats distribution.

We performed a mantel test to examine the correlation between genetic distance and geographic distance across individuals from Wales and England. The analysis revealed a positive statistically significant correlation (r = 0.333, p = 0.001), suggesting that genetic differentiation increases with geographic distance, demonstrating that individuals in Britain are isolated by distance.

Pairwise F_st_ values were computed to detect the genetic divergence between populations. Our results showed a comparison of English and Welsh populations having an F_st_ value of 0.036, compared to 0.106 between English and European and 0.145 between Welsh and European (Supplementary Figure S4).

### 3.3 Genomic Diversity in Britain in Comparison to Mainland Europe

As the British populations have undergone a severe population contraction prior to their current recovery, we investigated the impact of these demographic events on the genomic diversity of the different populations. European populations demonstrated a higher number of private SNPs (2,290,307) in comparison to English (838,471) and Welsh (578,869) (Figure 3a). This could be attributed to population structure and also demonstrates the genetic divergence between British and European mainland polecats. Welsh individuals showed a lower number of private SNPs in comparison to English individuals, although still demonstrating unique genomic diversity that still resides within the stronghold population.

**Figure 3.**
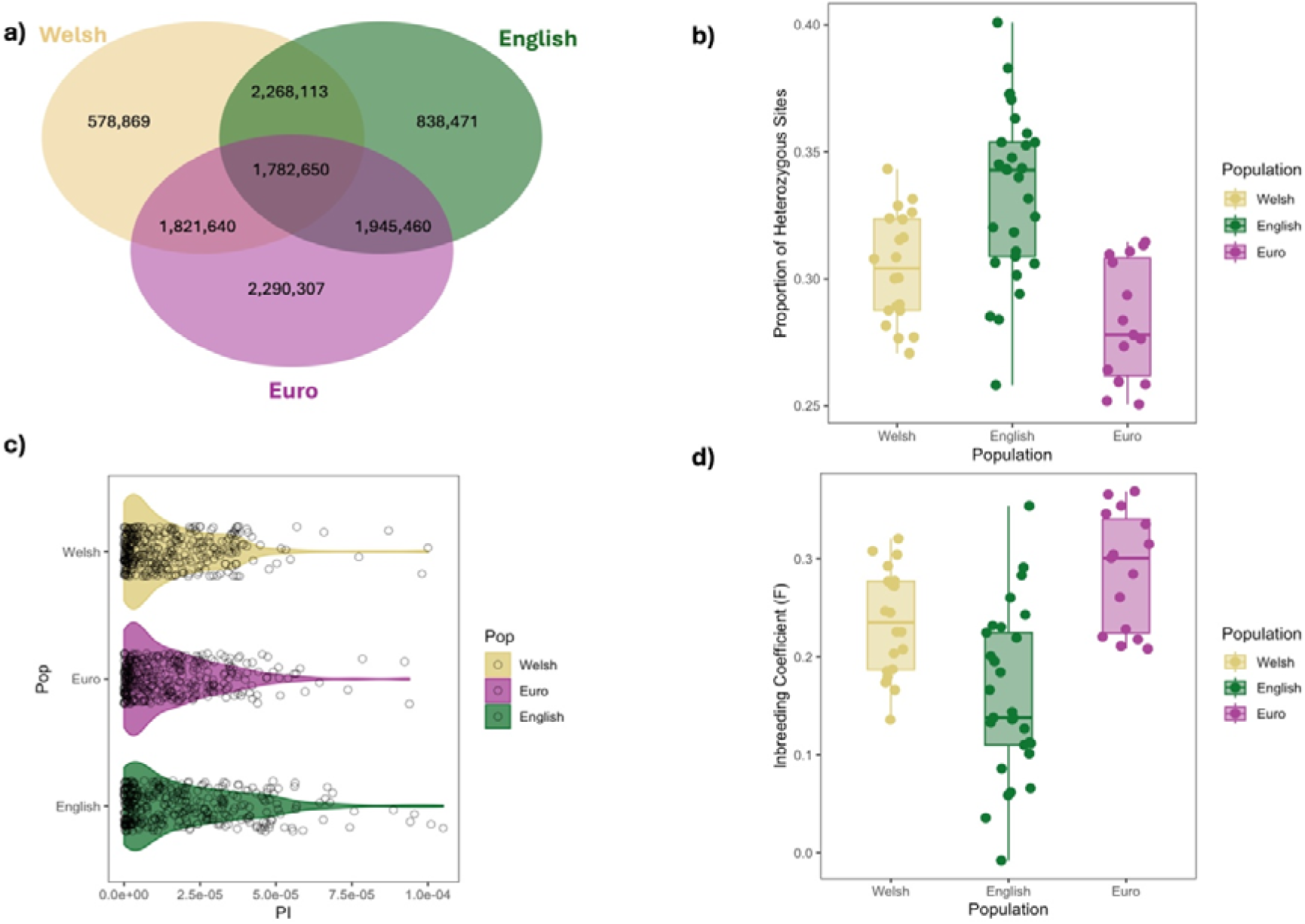
Genomic diversity estimates and comparisons across British and European populations (a) Shared and private SNPs between populations. (b) Proportion of heterozygous sites calculated using VCFtools (c) Population nucleotide diversity values (π), calculated in 20Mb windows across the genome and (d) Population-level inbreeding coefficient values

Nucleotide diversity estimates (Tajima’s π) (Supplementary table S4) revealed a significant difference amongst populations (H = 7.14, df = 2, p = 0.02), with the English population having significantly higher diversity when compared to the Welsh population (p <0.001) and European population (p = 0.01). However, there was no significant difference between the Welsh and European populations (Figure 3c).

Inbreeding coefficients (F2,60 = 17.6, p <0.001) and proportion of heterozygous sites (F2,61 = 18.16, p <0.001) significantly differed across the three populations (Figure 3b & 3d). Post-hoc comparisons found that the European population had a significantly higher inbreeding coefficient (p <0.001) and significantly lower proportion of heterozygous sites (p <0.001) in comparison to individuals in the English population. However, we did not account for population structure in the grouping of individuals due to the total sample size. French and Austrian populations had higher proportions of heterozygous sites and lower inbreeding coefficients than German, Italian, and Spanish populations (Supplementary Figure S7a & S7b). There was no significant difference between the Welsh and European populations for either inbreeding coefficient or the proportion of heterozygous site estimates. English individuals did show a significantly higher proportion of heterozygous sites (p <0.001) and significantly lower inbreeding coefficient (p <0.001) than those in the Welsh population.

### 3.4 Runs of Homozygosity (ROH) & Deleterious Mutations

To assess whether the Welsh population had undergone prolonged periods of inbreeding due to the population bottleneck, we investigated the presence of ROH within polecat genomes. We calculated the realised inbreeding coefficient (F_ROH_), the proportion of the genome that fell within ROH, which ranged from 0.33 - 0.76. Post-hoc comparisons showed that F_ROH_ values were significantly different between European and English individuals (W = 2.19, p = 0.0143) and English and Welsh (W = 2.42, p = 0.0077) individuals, but not between European and Welsh (W = 0.23, p = 0.4074) (Figure 4a & Supplementary table S5).

**Figure 4.**
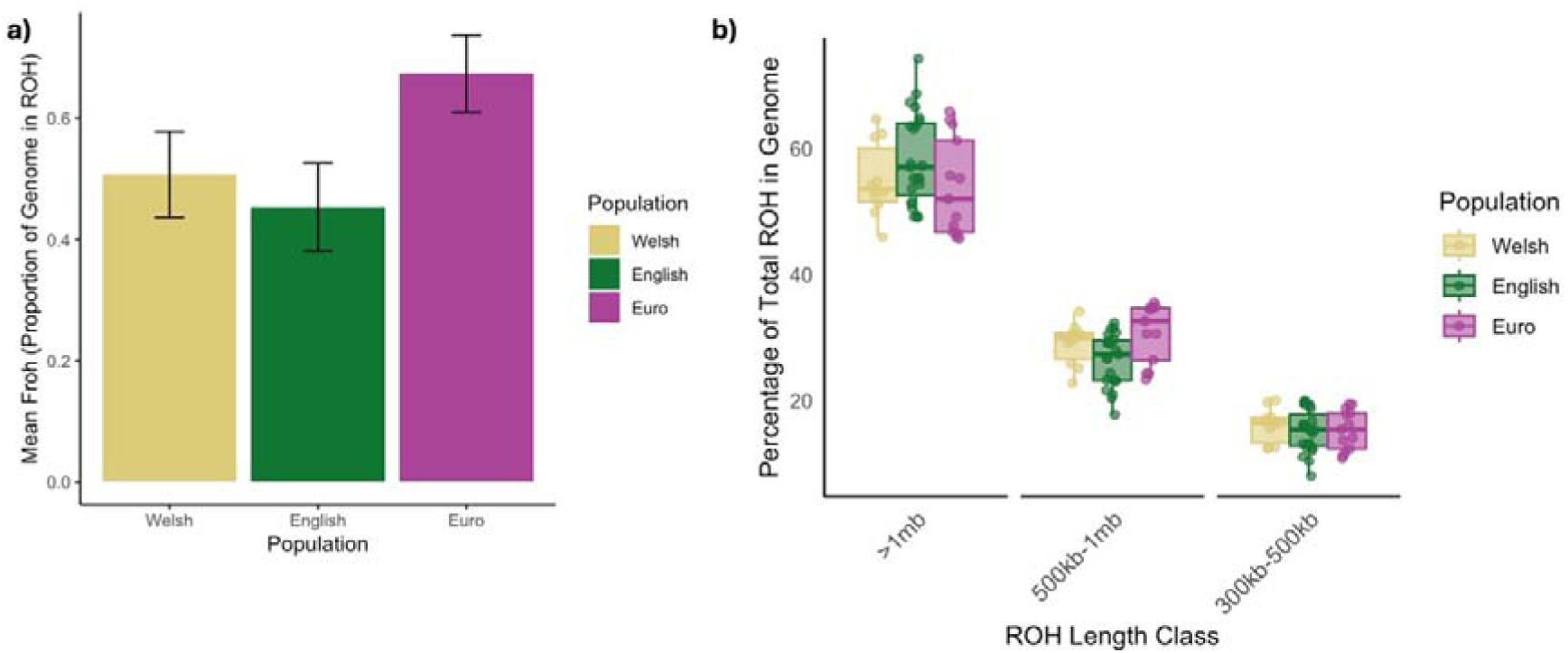
Population estimates of higher coverage individuals (mean depth >6x) and (a) the calculated proportion of the genome in ROH (F_ROH_) and (b) the distribution of ROH across different length categories. Data points represent the percentage of an individual’s total ROH that falls within each length category.

We separated ROH lengths into different classes to visualise short (300-500kb), medium (500-1Mb), and long (>1Mb) ROH across the genomes of individuals belonging to the Welsh, European, and English populations. Longer ROH are indicative of recent inbreeding events, with shorter ROH being indicative of historic events. Individuals belonging to the Welsh population were seen to have the lowest number of ROH segments in all length classes. Individuals belonging to the English population were seen to have a greater proportion of their genome within longer ROH segments (>1mb) (58.3% ± 1.50 SE) in comparison to the European (53.8% ± 2.14 SE) and Welsh (55.0% ± 1.90 SE) individuals (Figure 4b).

We investigated whether the mutation landscape differed among individuals belonging to European, Welsh, and English populations. We assessed derived mutational load across two functional categories as reported by SnpEff: missense and loss of function (LoF). We calculated both homozygous and heterozygous mutations per individual per category. Homozygous mutations in both categories were expected to be the most deleterious, and we also considered the load of heterozygous mutations.

Homozygous LoF mutation load differed significantly between populations (F2,60 = 16.81, p <0.01), with both Welsh and English populations showing a significantly higher homozygous load within LoF mutations in comparison to the European populations (p <0.01) (Figure 5a). There was also a significant difference amongst populations in LoF masked load (F2,60 = 5.584, p <0.01), with individuals in the English population having a significantly higher masked load than Welsh individuals (p <0.01).

**Figure 5.**
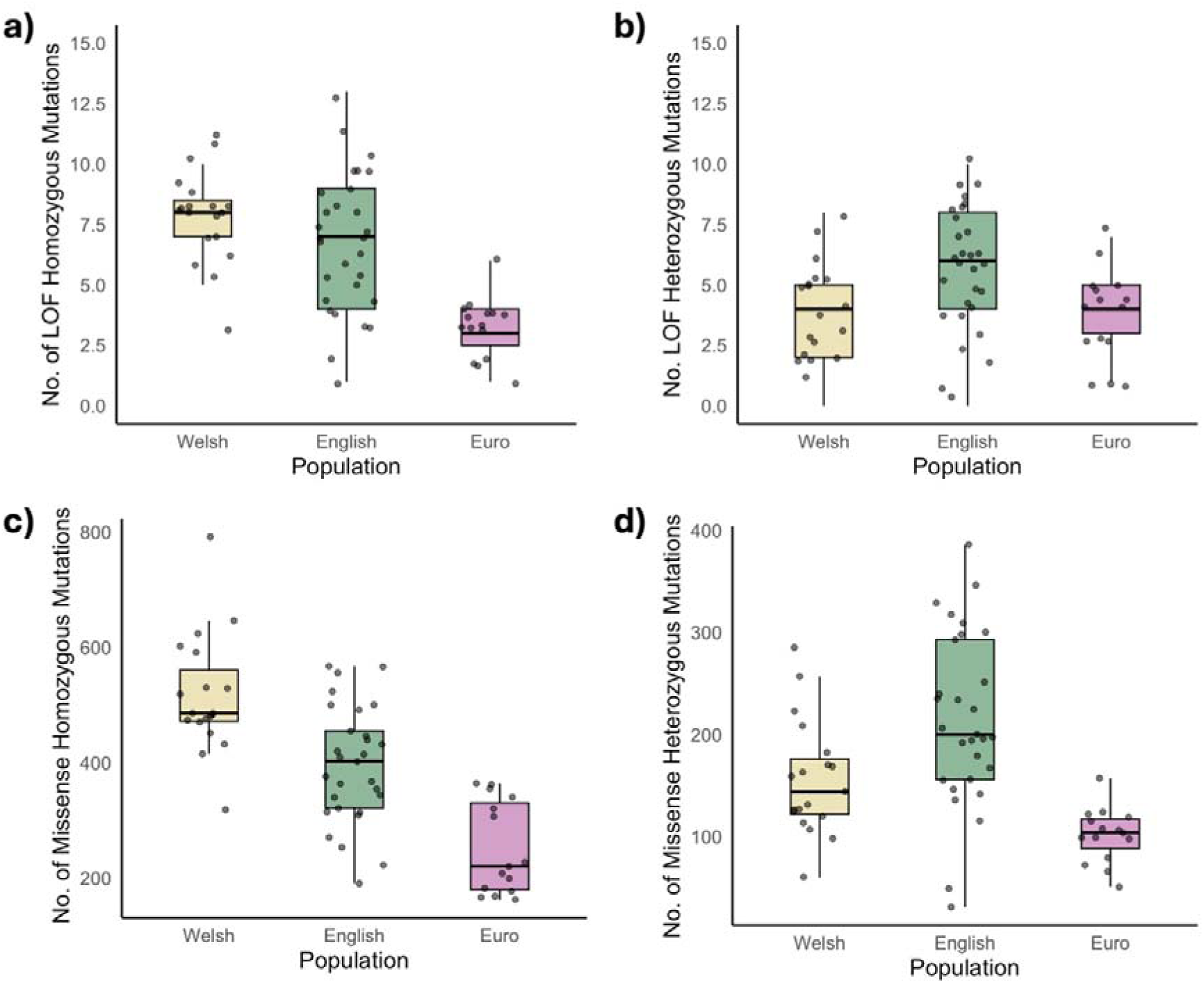
Mutation load landscape across populations within different functional categories. Loss of function (LoF) (a) homozygous mutational load per population and (b) heterozygous mutational load and missense (c) homozygous mutational load and (d) heterozygous mutational load.

Missense homozygous mutations significantly differed between populations (F2,60 = 31.39, p < 0.01). Post-hoc comparisons revealed that Welsh individuals had significantly more missense homozygous mutations than English and European individuals (p <0.01) (Figure 5c).

Masked, heterozygous missense load also significantly differed among populations, (F2,60 = 14.77, p < 0.01), with English individuals showing a higher masked load than both Welsh (p <0.01) and European individuals (p <0.01) (Figure 5d).

### 3.4 Historic effective population size (N_e_)

We used GONE to estimate the recent historic effective population size for the British population of European polecats. Our estimates using GONE correlate with a demographic decline in population size occurring around 30-40 generations ago - which using an estimate for generation time of the polecat of 4 years (Di Marco et al. 2013), corresponds to the mid-late 19th century (1854-1894) (Figure 6). Recovery of the polecat population in Wales based on GONE estimates fits with previous estimates of 10-12 generations ago (1966-1974) (Supplementary table S7).

**Figure 6.**
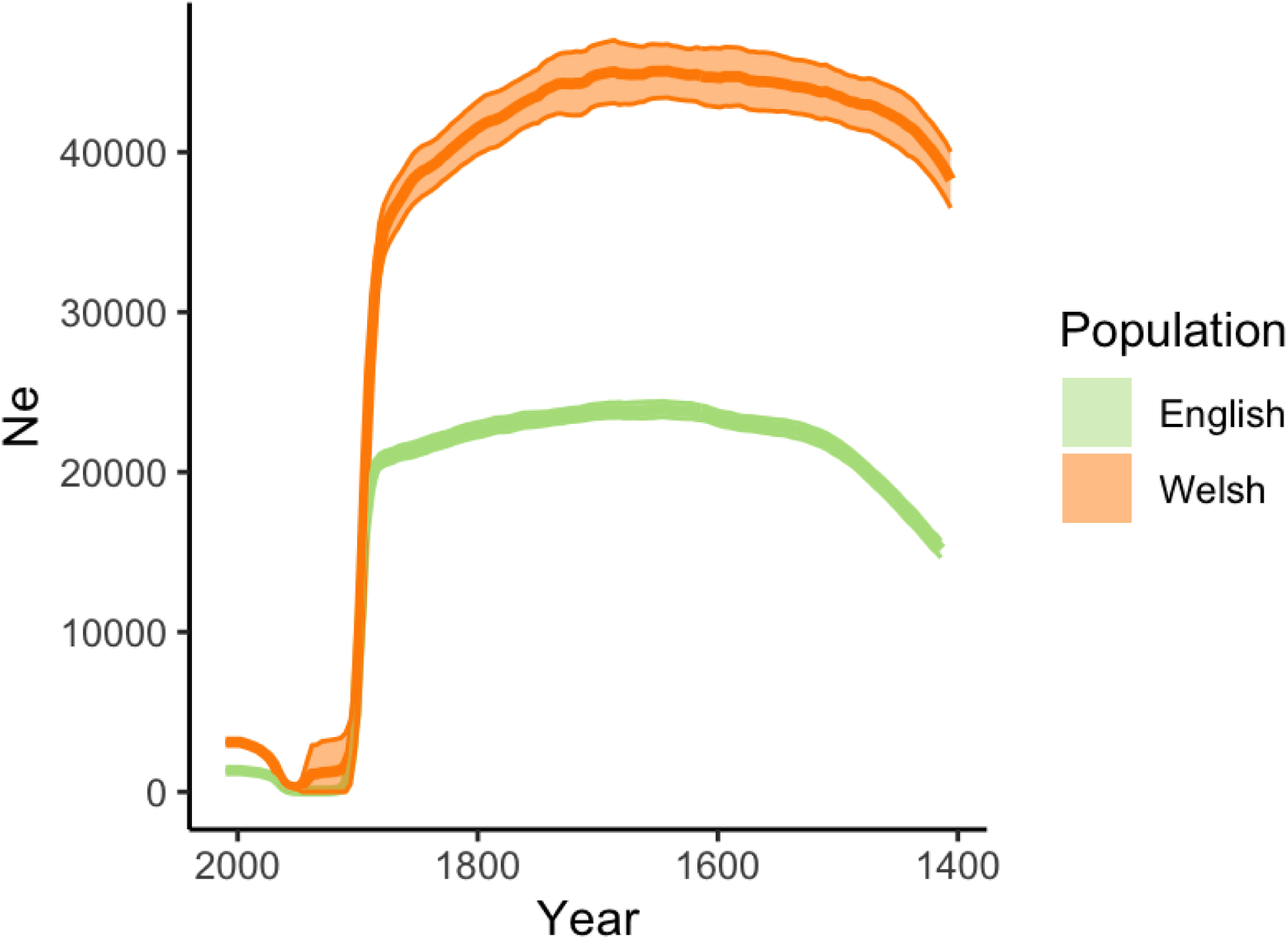
Estimated mean effective population size (solid line) with 95% confidence intervals (shaded area) for English (green) and Welsh (orange) populations across 200 generations (generation time = 4 years) based on whole genome SNP data computed in GONE.

## 4.0 Discussion

Here, we provide the first evidence of a genetic bottleneck having occurred within the British population of polecats. Our analyses indicate that the population drastically decreased in effective population size from the late 1800s, with this bottleneck persisting for ca. 100 years. Our analyses also indicate that the population began recovering from 1960-70 onwards. This coincides with a prolonged period of relaxed persecution, and the introduction of protective legislation. Our results demonstrate that the British polecat population is distinct from European mainland populations, and is recovering from the genetic bottleneck.

Conservation efforts should focus on increased protection measures and monitoring of the Welsh population, as it represents unique genetic integrity in comparison to admixed, English populations. Our study also highlights the importance of genomic data in being able to detect signatures within populations to fully detect and monitor the success of their recovery.

### 4.1 Comparison of British to Mainland European populations

Our analyses indicate that the British polecat population is genetically diverged from European mainland populations. This is highlighted in pairwise F_st_ estimates per population where a higher F_st_ value (closer to 1.0) suggests a population has low levels of breeding and therefore is more genetically distinct from populations compared to lower F_st_ values. Our results indicate that English and Welsh populations have a low fixation index (0.03) and are therefore likely to share genetic variation. F_st_ values between English and European (0.106) and European and Welsh populations (0.145) are slightly higher, suggesting a greater degree of genetic distinction between mainland European and British polecats, which is also demonstrated by the degree of private and shared polymorphisms across the populations (Figure 3a).

This is also evident from our ADMIXTURE analyses which further corroborates the genetic divergence seen between Welsh and European polecats (Etherington et al. 2022). British and European polecats are thought to have been separated for ca. 8ky (Montgomery et al. 2014), since the flooding of the Doggerland bridge after the Last Glacial Maximum. Several widespread mammal populations that are native to Britain demonstrate genetic distinction due to isolation over time from mainland European populations. For example, Irish and British populations of red fox (*Vulpes vulpes*) demonstrate unique genetic diversity that distinguishes them from mainland European populations (McDevitt et al. 2022). Likewise, native British populations of roe deer (*Capreolous capreolus*) demonstrate genetic differentiation from mainland European populations, most likely due to colonisation and isolation events (de Jong et al. 2020). Populations of British red deer (*Cervus elaphus*) also demonstrate genetic distinction from mainland European populations, but show a slightly lower genetic distance to Iberian populations, potentially suggesting retention of genetic diversity within populations from similar migration waves during postglacial colonisation (Carranza et al. 2024). Therefore, the genetic distinction we observe between Welsh and European mainland polecats likely results from their recent isolation, mirroring the pattern of genetic divergence seen across other British populations of wide-ranging mammals (Montgomery et al. 2014; McDevitt et al. 2022; Carranza et al. 2024).

### 4.2 Population Structure

Our analyses indicate population structuring based on geographic location within polecats in Britain (Figure 2b). Ancestry estimates also demonstrate a higher incidence of introgression in polecats that are found at the furthest edges of the polecat range from the Welsh refugium population (Figure 2c). We also observe a similar relationship in our maximum likelihood phylogeny, whereby English individuals that are found at the more eastern and southern edge of the polecat’s range are clustered more closely to domestic ferrets than to the Welsh individuals (Supplementary Figure S6). In agreement with (Costa et al. 2013), we find the least introgression of domestic ferret genes within individuals found in Wales and English border counties. This observed pattern fits the narrative that as the polecat population expanded eastwards, individuals hybridised with feral domestic ferrets within those regions. However, the presence of introgressed polecats within these regions may also have been exacerbated by covert reintroductions of individuals from hybrid stocks (Birks and Kitchener 1999).

### 4.3 Demographic Analyses

Using whole genome sequence data, it was possible to detect and analyse historic trends in effective population size in the British population of the European polecat. The British polecat bottleneck was thought to have occurred between the late 1800s to the early 1900s, with populations showing recovery since the 1970s (Langley and Yalden 1977). The most recent historic genetic bottleneck was evident in our results using GONE, with a sharp decrease in N_e_ occurring in the Welsh population around 30-40 generations (120-160 years ago), from the sampling time of 2014-2015. This coincides with the known history of persecution of polecats from gamekeepers, as with other populations of terrestrial carnivorans in Britain, which subsided for a period during and following World War I, and again after legal protections came into place (Langley and Yalden 1977; Sainsbury et al. 2019). Alongside legal protections, it is thought that increasing rabbit (*Oryctolagus cuniculus*) populations, an important prey for polecats (Sainsbury et al. 2020), which had declined due to myxomatosis and began recovering from the mid-20th century, are also closely linked with the recovery of the polecat (Langley and Yalden 1977; Sumption and Flowerdew 1985; Sainsbury et al. 2019).

We observed that historic N_e_ estimates were lower for the English population than the Welsh (Figure 6, Supplementary table S8). This could be explained both by the rapid expansion of individuals from Wales to England and by the hybridisation and subsequent introgression of those individuals with populations of feral domestic ferrets. Our results also highlight that modelling demographic histories can be impacted by hybridisation and population structure (Novo et al. 2023). Previous work (Costa et al. 2013), using microsatellites in British polecats could not detect any evidence of a genetic bottleneck. This could be due to the lower resolution from the smaller representation of genetic markers used but could also be due to the admixture from domestic ferret populations as suggested by the authors. We observe this pattern within the English population having greater levels of introgression than the Welsh population and therefore observe not only a lower historic effective population size but also a lower present day effective population size when just modelled using the English population. We see that population structure can obscure the recent historic effective population size when trying to model the European population’s demographic histories (Supplementary Figure S8). Modelling these as a whole population is difficult due to the different environmental pressures that each population faces. GONE authors advise not to model demographic histories with less than 5 individuals (Novo et al. 2023), of which only the Italian population was viable from our European samples. However, our model reconstruction demonstrated the unreliability of this with low power and possible population structure (Supplementary Figure S8).

Census data in 2018 estimated that the polecat population in Britain was currently 88,300 (Mathews et al. 2018). Our estimates corroborate survey data from the Vincent Wildlife Trust that the Welsh population is expanding, with more ‘true polecat’ sightings in Welsh counties and border counties such as Herefordshire and Shropshire (Croose 2016), which are areas that were thought to contribute to the historic stronghold of the polecat (Langley and Yalden 1977), and these areas also having the individuals with the least admixture (Figure 2c).

### 4.4 Genomic signatures associated with the population bottleneck

We hypothesised that we would see genomic signatures within the population associated with the prolonged genetic bottleneck period. Our results demonstrate the presence of potentially harmful deleterious mutations within both Welsh and English individuals. It also shows that heterozygous, potentially masked deleterious variants are more prevalent in admixed English individuals, and that the Welsh population has a higher unmasked load of missense variants. This is as expected, as periods of inbreeding in Welsh populations would lead to a higher presence of homozygous deleterious variants within the population. English individuals appear to have higher heterozygosity, due to the introgression with domestic ferret. This is also confirmed in the higher nucleotide diversity across the genomes of English individuals. However, no significant difference amongst populations was found in mutations that were categorised as homozygous LoF, suggesting these mutations may have been purged from the Welsh population due to their expansion and purifying selection acting upon them. Further investigation into historic genomic signatures through the addition of historical samples would provide an insight into temporal changes of mutational load over time, which would assist in resolving potential signatures of purifying selection within the Welsh population, and would be consistent with other studies that have observed this (van der Valk et al. 2019; Grossen et al. 2020; Dussex et al. 2021)

Individuals belonging to the Welsh population had higher overall ROH in their genomes in comparison to the English population (Figure 4a). This could be due to the long-term, low effective population size within the Welsh refugium. Welsh individuals also demonstrated a higher mean of shorter ROH within the genome in comparison to the English populations. This is indicative of more ancestral inbreeding events, and is a strong indicator that the population underwent a bottleneck further back in time (Saremi et al. 2019). Longer ROH lengths can indicate recent inbreeding events, as recombination has not yet acted on breaking apart these segments (Ceballos et al. 2018). Individuals belonging to the English population were shown to have a higher mean number of long ROH in their genome than Welsh. This could be associated to the introgression with feral domestic ferrets that have been subjected to years of selective, human-mediated breeding. However, it may also be indicative of the historic bottleneck, followed by the expansion of some individuals to low areas of genetic connectivity and therefore increased occurrence of inbreeding events ((Manunza et al. 2016; Colpitts, McLoughlin, and Poissant 2022).

European individuals overall showed a higher mean F_ROH_ than both English and Welsh individuals, as well as a higher mean proportion of medium length ROH. However, across the length classes and F_ROH_ values there was a varied ROH landscape per population (Supplementary Figure 7c). This could be due to population-level pressures persisting in their environment at the time (Croose et al. 2018) and may indicate the necessity for similar genomic studies in polecat populations in regions of Europe where the population is declining.

### 4.5 Conservation Implications

Here, we provide the first evidence of the severity of the population bottleneck experienced by the Welsh polecat population from the late 1800s to the mid-1950s. We demonstrate that the population started to expand when persecution pressures were lessened and after protective legislation came into action, thus supporting legal protection as a conservation management tool.

We present a unique scenario of population recovery in a once heavily persecuted mammal in Britain. The recovery can largely be attributed to protective legislation and the resurgence of rabbit populations following the myxomatosis outbreak, as well as to the introgression with feral domestic ferrets, which has facilitated the eastern range expansion of the population. Genomic introgression with a domesticated species is not a typical conservation management tool but has been witnessed to assist in the recovery of other species, such as the American bison (*Bison bison*) (Stroupe et al. 2022), and Iberian ibex (*Capra pyrenaica*) (Münger et al. 2024). However, the decline of the wildcat (*Felis silvestris*) population in Scotland has been severely hampered by hybridisation and subsequent introgression with the domestic cat (Howard-McCombe et al. 2023). It could be that due to the relatively recent nature of the hybridisation of the European polecat and domestic ferret in Britain, the harmful consequences are yet to surface. However, (Costa et al. 2013) demonstrated that the lack of F1 hybrids observed within the British population suggests that these hybridisation events are rare. Feral ferret populations are hard to distinguish in mainland Britain but are known to thrive on surrounding islands where there is a lack of or reduction in competition from polecats and other predatory mammals (Kitchener and Birks 2008). This indicates that F1 hybridisation events between feral ferrets and polecats in wild populations are likely to be rare, yet they still represent a potential threat to the genetic integrity of the recovering polecat population.

Our analyses also demonstrate that the Welsh population holds unique genomic diversity that could be important for the resilience of the population. This could have important conservation implications for reintroducing polecats from Wales to other parts of the country. Additionally, conservation concerns are underscored by the unmasked missense mutational load persisting in the genome as a result of the prolonged low effective population size (Robinson et al. 2023). The monitoring of this elusive species is hard, due to its low population density and nocturnal habits (Croose et al. 2018; Barrientos et al. 2024). Monitoring through non-invasive methods, such as the Vincent Wildlife Trust survey, where sightings and suspected roadkill are reported, has proven successful and has helped to document the polecat’s recovery within Wales and England (S. 2008; Croose 2016). However, these efforts are not matched in some parts of mainland Europe and therefore the polecat’s status in some countries needs to be clarified (Kominos and Galanaki 2018; Hofmeester et al. 2019).

In a study aiming to document the status of the polecat across its known range, of the 34 countries where data were found, 20 countries reported that the polecat population was declining (Croose et al. 2018). Only two countries reported the polecat population as increasing, i.e., Estonia and Britain, although there is some evidence that populations in Poland could be genetically stable (Martínez-Cruz, Zalewska, and Zalewski 2022). Therefore, conservation priorities should be to continue monitoring the Welsh population to protect its genetic legacy and ensure the population’s continued expansion. Given the reported declines in other known polecat ranges (Baghli and Verhagen 2003; Costa, Fernandes, and Santos-Reis 2014; Barrientos 2015; Szatmári et al. 2021; Frantz et al. 2022), conserving the Welsh population is crucial for supporting the global population. British polecats face threats not only from hybridisation with feral ferrets, but also from road traffic collisions, which hinder movement during peak dispersal (Barg, MacPherson, and Caravaggi 2022). Conservation efforts should therefore prioritise reducing habitat fragmentation and minimising the need for polecats to cross major roads.

It should also be noted that the genetic recovery of a population can often lag behind its demographic recovery (Adams and Edmands 2023; Thomas et al. 2022; Dussex et al. 2023). In the case of the British polecat population, we do see genetic structuring due to admixture across parts of England. Another British mustelid, the Eurasian Otter (*Lutra lutra*), has subsequently undergone a recent historic genetic bottleneck and is recovering across its previous ranges (Thomas et al. 2022; du Plessis et al. 2023). However, barriers, such as landscape changes, have potentially contributed to the fragmentation and low genetic connectivity between populations. We recommend that continued monitoring be complemented with continued analysis of genetic data, to ensure that polecat populations in Wales do not become genetically isolated due to landscape change and continue to recover both demographically and genetically (Thomas et al. 2022).

This study examined the recovery of polecats across much of Great Britain, however it did not include data on the population in Scotland. According to survey records, it was thought that by 1915, polecat populations were eradicated throughout Scotland. Covert re-introductions of polecat were thought to have occurred across some areas in southern England, Cumbria and Argyll (Davison et al. 1999; Birks and Kitchener 1999). Previous work has highlighted the more ‘ferret-like’ ancestry of polecats found within regions of Scotland (Costa et al. 2013), with feral ferret populations being found on offshore Scottish islands. The 2014-15 survey conducted by the Vincent Wildlife Trust recorded only 29 sightings of polecats, representing just 2% of the total records from Britain (Croose 2016). However, some of these sightings occurred in a newly identified distribution area in Dumfries, which may indicate a range expansion of the Cumbrian populations. Further sampling and genomic analyses, in addition to the continued monitoring efforts, would provide greater insights into the population structure and conservation status of the remaining polecat population within Scotland.

## Acknowledgments

RS was supported by a NERC PhD studentship through the ARIES DTP (NE/S007334/1, 2465312). WH and GE acknowledge the support of the Biotechnology and Biological Sciences Research Council (BBSRC), part of UK Research and Innovation; Earlham Institute Strategic Programme Grants Cellular Genomics BBX011070/1, Decoding Biodiversity BBX011089/1, Delivering Sustainable Wheat BBX011003/1 and their constituent work packages (BBS/E/ER/230001A, BBS/E/ER/230001B, BBS/E/ER/230001C, BBS/E/ER/230002A, BBS/E/ER/230002B, BBS/E/ER/230003A, BBS/E/ER/230003C). The author(s) acknowledge support from the Biotechnology and Biological Sciences Research Council (BBSRC), part of UK Research and Innovation, Core Capability Grant BB/CCG1720/1 supporting the work delivered via the Scientific Computing group, as well as support for the physical HPC infrastructure and data centre delivered via the NBI Computing infrastructure for Science (CiS) group.and the National Capability BBS/E/T/000PR9816. ACK thanks the Negaunee Foundation for its continued support of a curatorial preparator who sampled many of the polecats used in this study. We are also grateful to Federica Di Palma for her early input to the project.

## Author Contribution

The language used to recognise author contributions is described by CRediT Taxonomy (https://credit.niso.org/).

Conceptualization: GE, WH; formal analysis: RS; funding acquisition: GE, WH, FDP; investigation: RS; methodology: RS, GE, WH; project administration: RS, GE, WH; resources: JM, ACK, GE, WH; software: RS; supervision: JM, GE, WH; visualisation: RS; writing - original draft: RS; writing review & editing: RS, JM, ACK, GE, WH

## Data Availability

All raw reads for the sequences generated during this study have been uploaded to ENA with the project number PRJEB76468. Bioinformatic scripts used to carry out analyses can be found at this repository (https://github.com/TGAC/EuropeanPolecatPopGen/).

## Supplementary material

Supplementary methods, figures and tables are available in the links provided.

