## Supplementary figures for "Characterisation of the historic demographic decline of the British European polecat population"

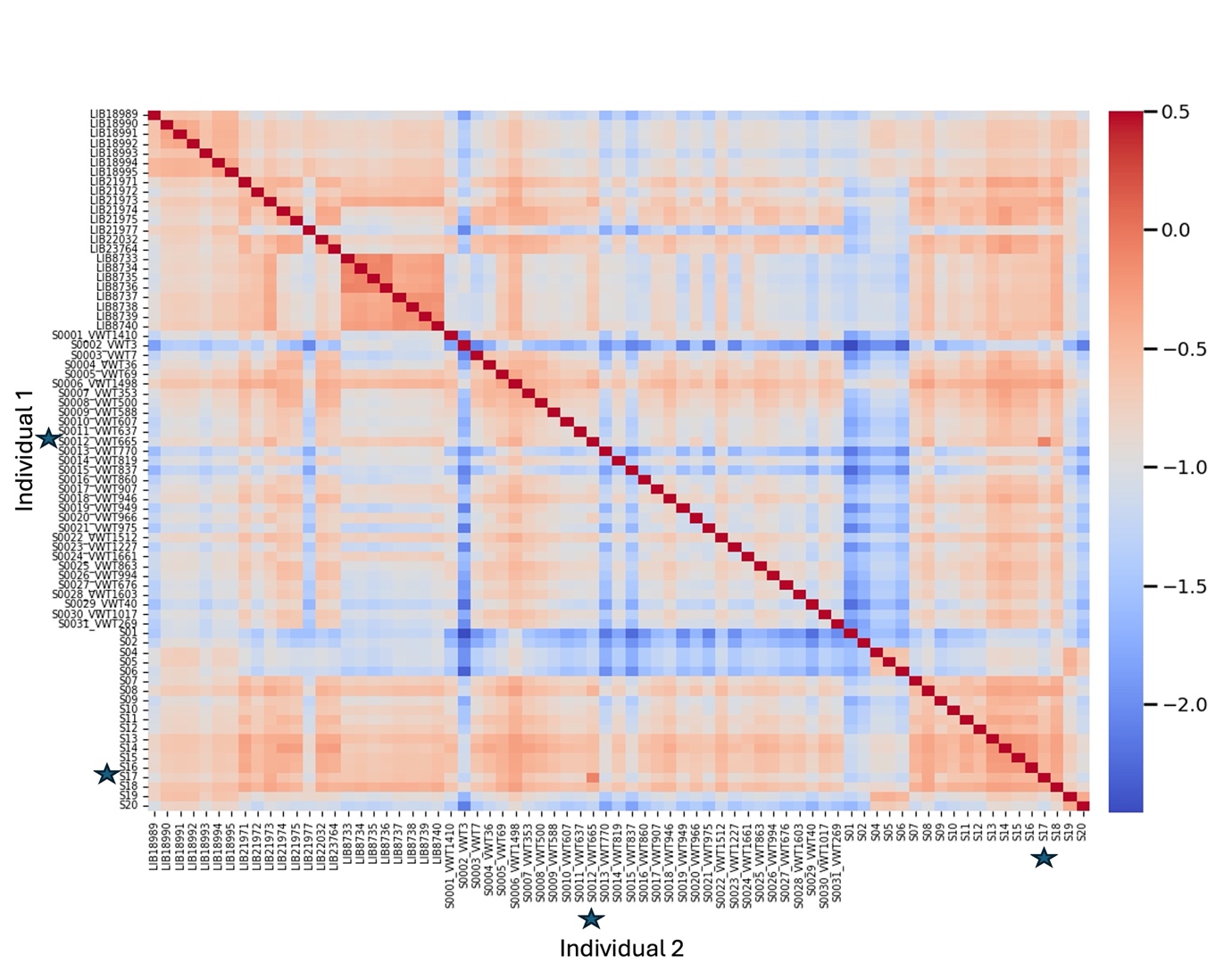

**Figure S1**: Kinship coefficients between each of the 65 polecat and 8 domestic ferret individuals. KING estimates a relatedness-PHI score which can range from >0.354, [0.177, 0.354], [0.0884, 0.177] and [0.0442, 0.0884] and corresponds to duplicate/MZ twin, 1st-degree, 2nd-degree, and 3rd-degree relationships respectively. Starred individuals (S17 & S0012_VWT665) were identified as having a 3rd-degree relationship (0.076) and S17 was removed from further analyses.

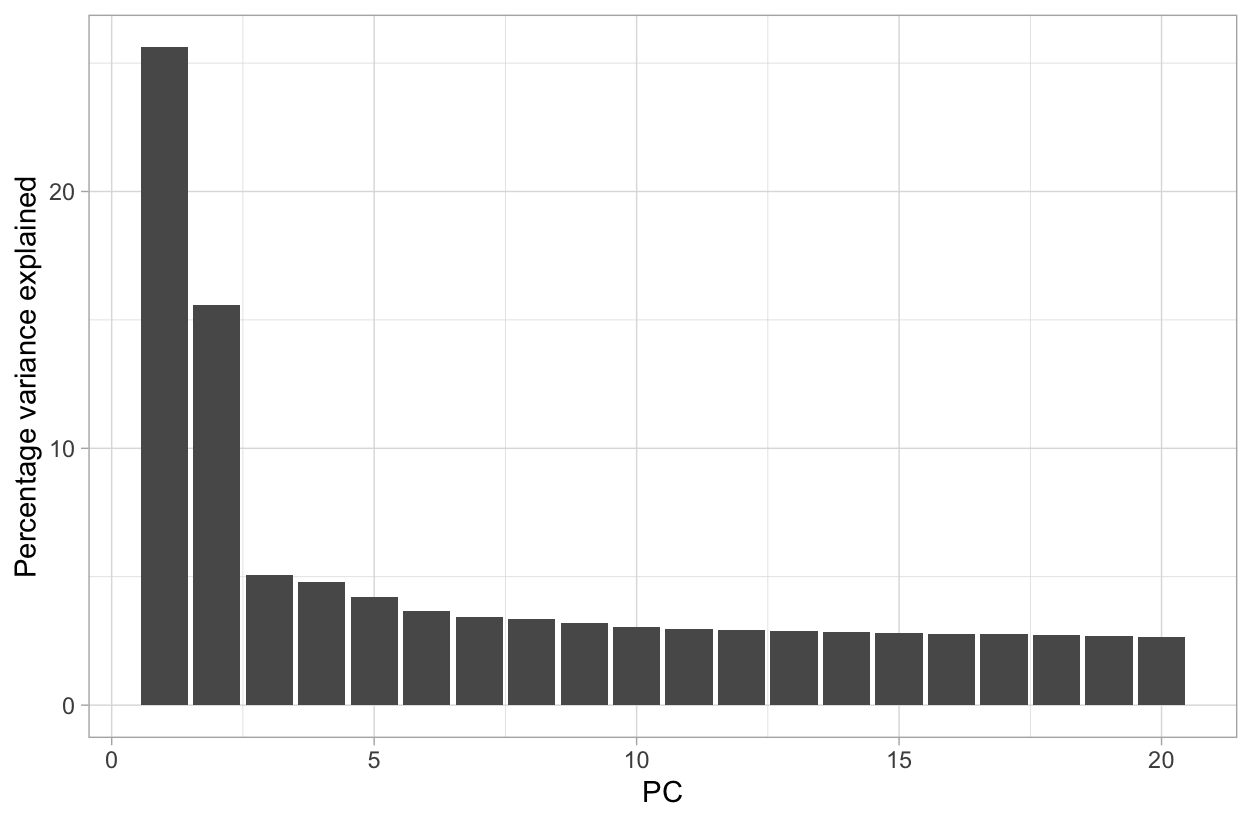

**Figure S2**: Percentage of variance explained by PC’s 1-20 using whole-genome SNPs.

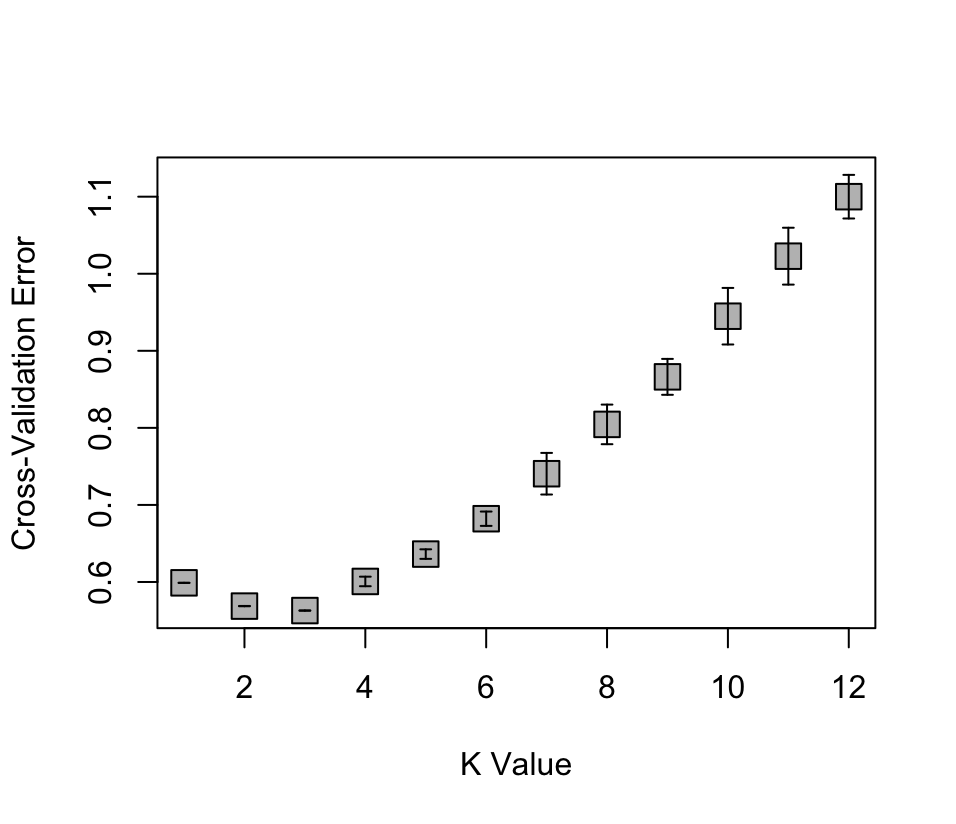

**Figure S3:** Average K value for replicated runs of ancestry for K 1-12**.**

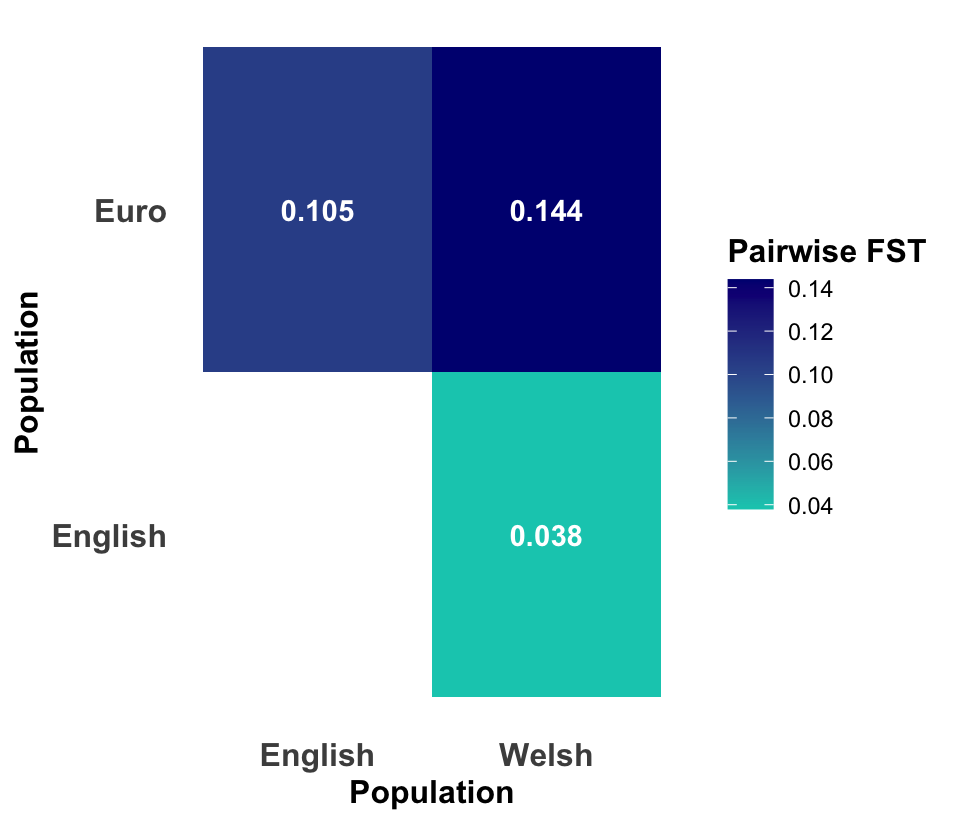

**Figure S4**: Pairwise F_ST_  values for whole genome SNPs between populations.

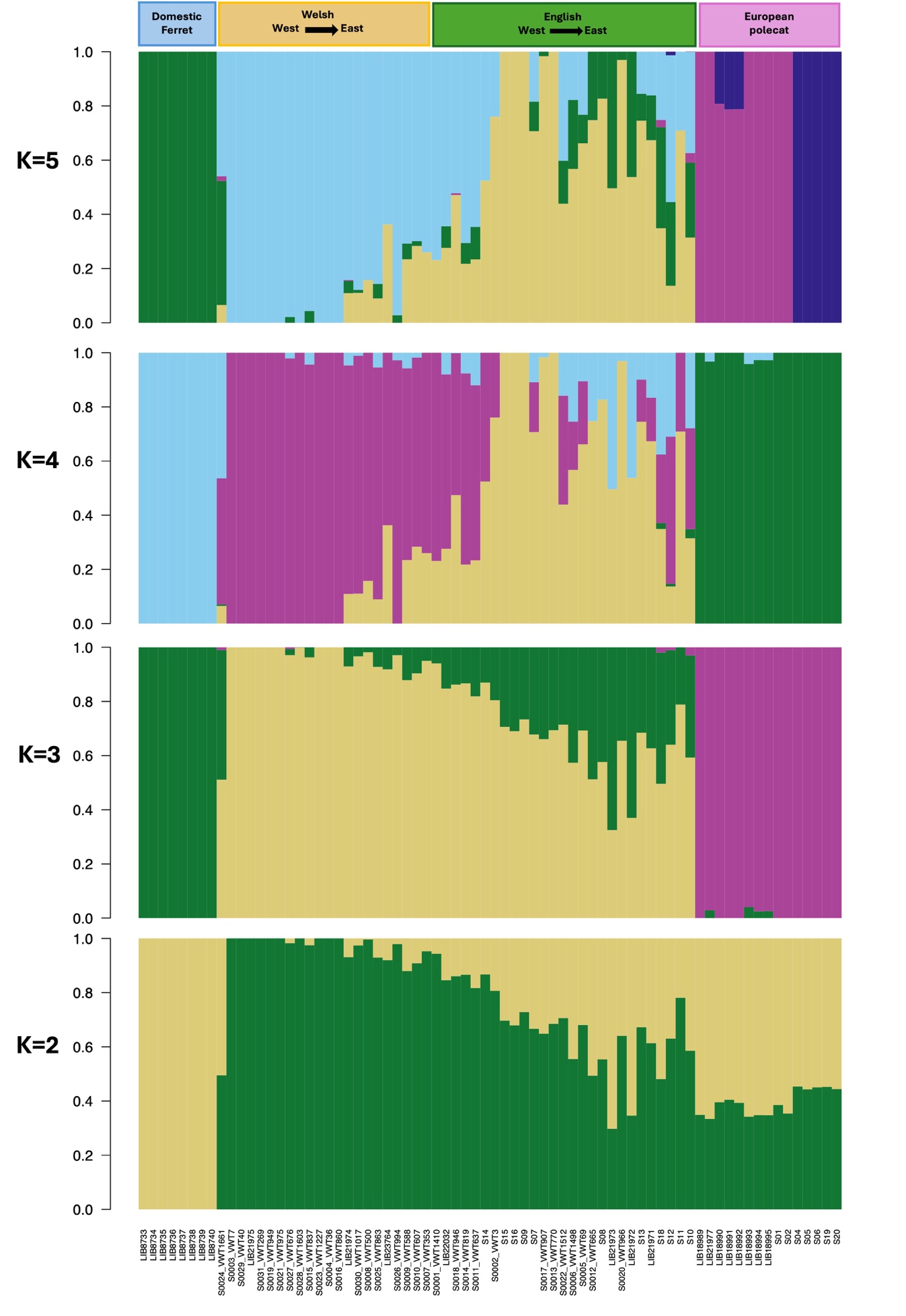
**Figure S5:** Ancestry proportions for each individual inferred from ADMIXTURE for K values of 2-5.

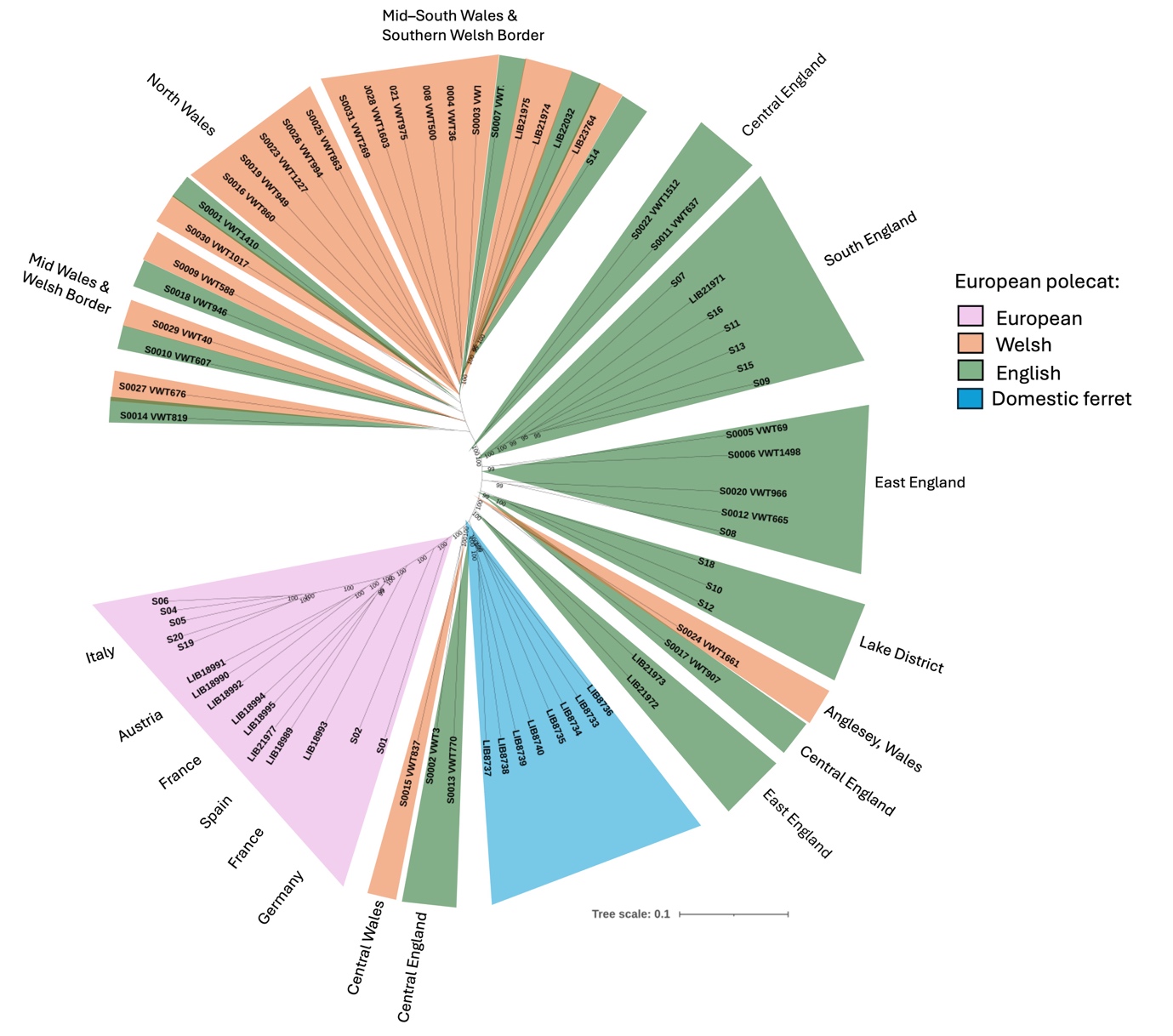

**Figure S6**: Maximum likelihood phylogeny with 100 bootstrap replicates for genome-wide SNPs from 64 samples. The number at nodes refer to bootstrap values of 95 and above. Branch lengths are in expected substitutions per site.

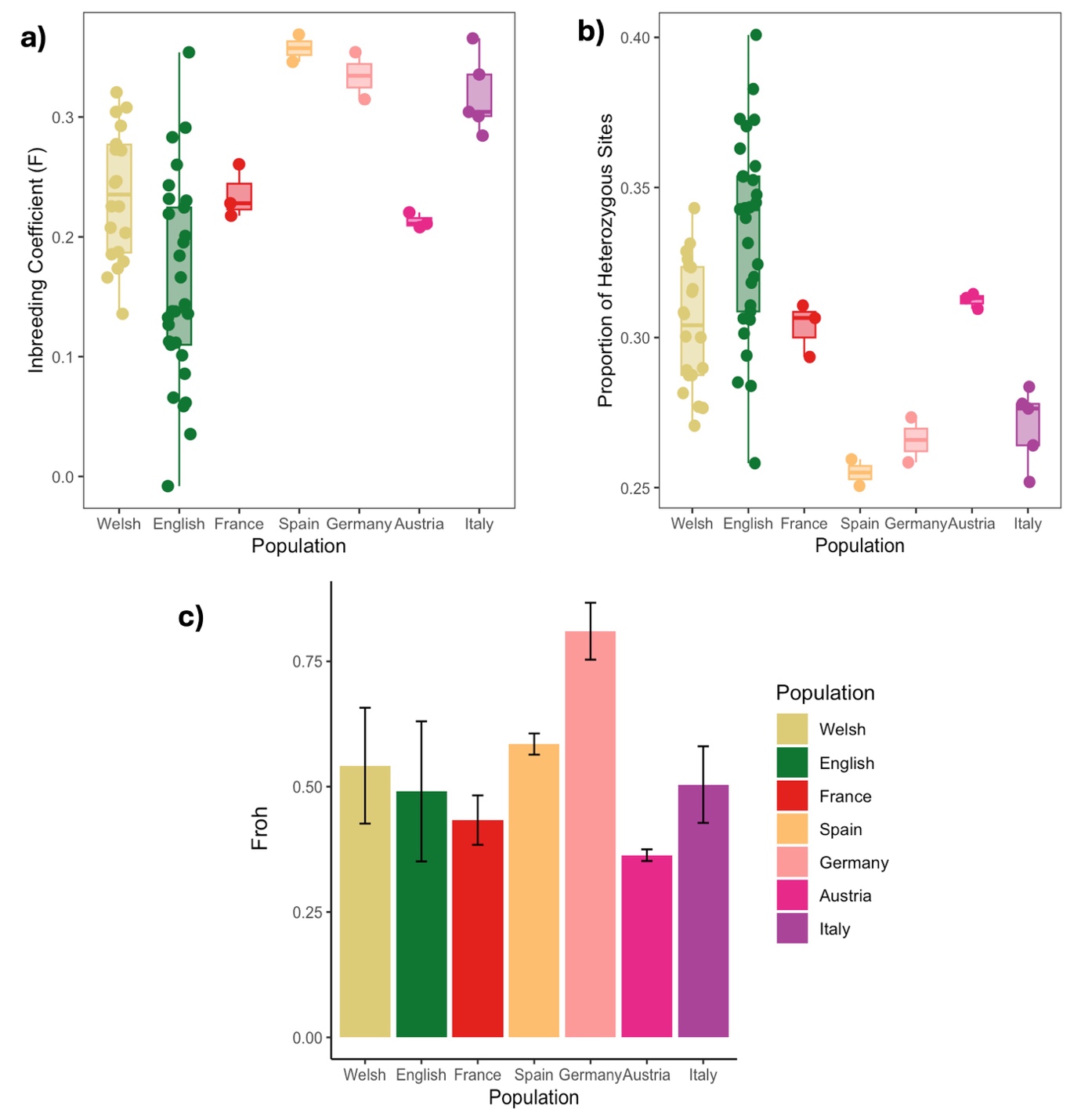

**Figure S7**: Inbreeding estimates and heterozygosity comparisons across British and European populations (a) Population-level inbreeding coefficient values (b) Proportion of heterozygous sites calculated using VCFtools and (c) Mean F_ROH_ per population

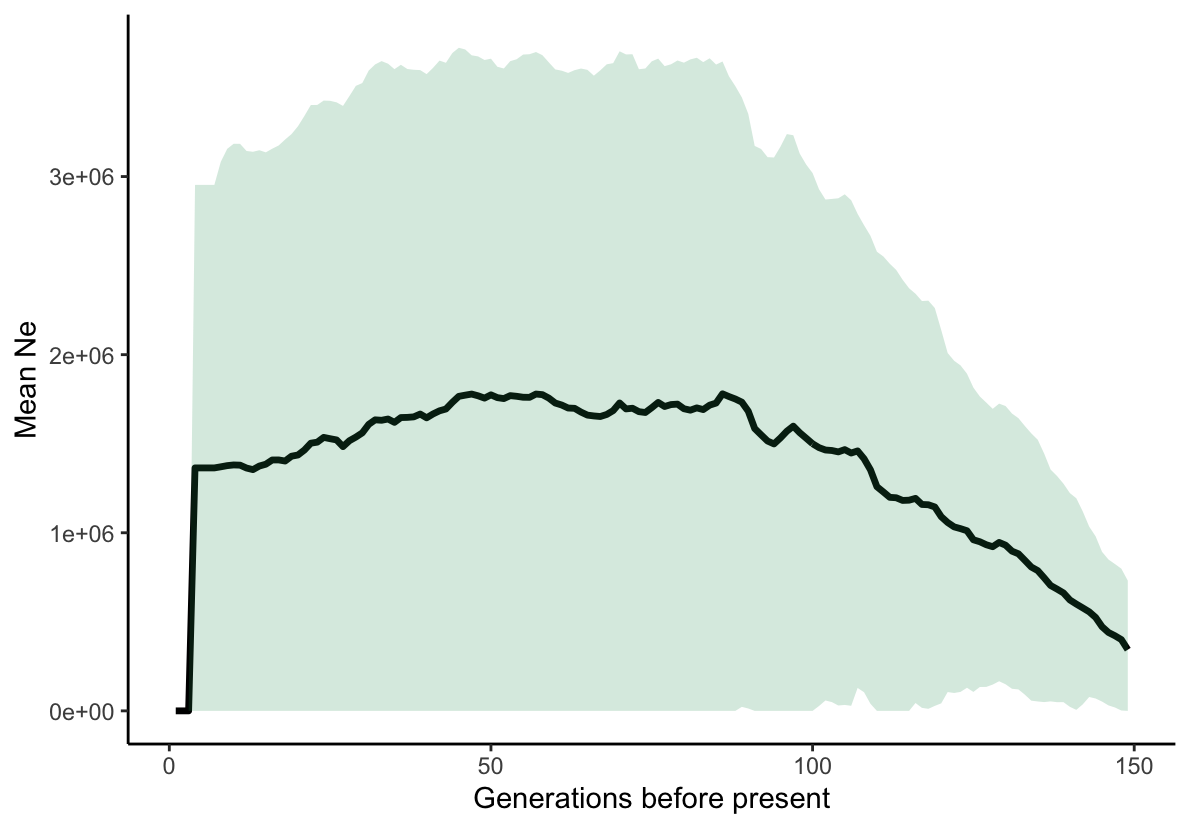

**Figure S8:** Estimated mean effective population size (solid line) with 95% confidence intervals (shaded area) for the Italian polecat population across 200 generations (generations = 4 years) based on whole genome SNP data computed in GONE.
