## Supplementary methods for "Characterisation of the historic demographic decline of the British European polecat population"

**Alleviation of batch effect from samples**

Data curated for this study were produced from different sequencing experiments and had been generated using different library preparation methods, run on different sequencing technology platforms and varied in sequence length and coverage (Supplementary table 1).

The first attempt at calling variants was done so using bcftools mpileup. Figure 1a and 3a demonstrate the PC’s run using these variant datasets, demonstrating the issue of batch effect arising within our data. To tackle the impact of batch effect on the real biological signal (Leek et al. 2010) and consistent with previous studies (Leigh et al. 2018), we trimmed down sequence read lengths so that all reads were of similar sizes (100bp) using TRIMMOMATIC (version 0.39) (Bolger, Lohse, and Usadel 2014). We then re-aligned the reads to the reference genome with BWA MEM (version 0.7.17) (Li and Durbin 2009), and followed the GATK version 4.5.0 (McKenna et al. 2010) pipeline for variant calling. Figure 2a demonstrates the PC’s run using this variant dataset, where we see an improvement in the resolution between samples and technology.

Previous work by (Lou and Therkildsen 2022) demonstrated that downsampling and base recalibration of quality scores on datasets from different sequencing runs could successfully alleviate batch effects. Therefore, we carried out a further alleviation step of downsampling samples to a similar mean (~5x) sequencing depth using samtools. Briefly, we calculated proportions (Supplementary material S1) to downsample reads within bam files by to reduce the average read coverage to be consistent with other samples We then used the BaseRecalibator approach implemented by the GATK pipeline based on their best practice data pre-processing and best practice workflow for SNP discovery (van der Auwera and O’Connor 2020).Variants were then called per sample, before being consolidated into one VCF for all samples. These were then filtered to include only SNPs and based on quality score.

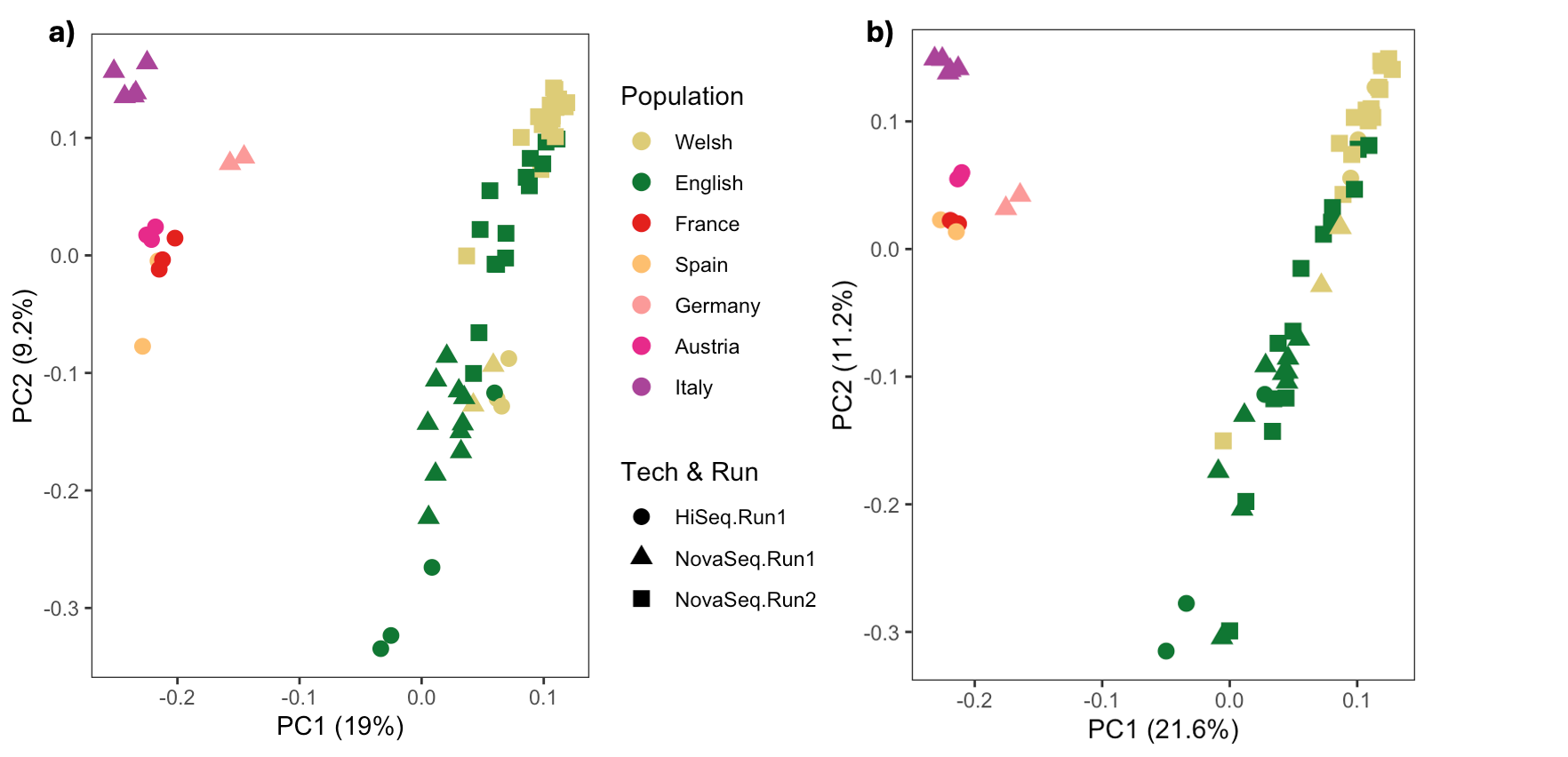

.

Figure 1: Principal component analysis (PCA) of whole genome SNP’s for British and European samples. Figure **a)**  Without being batch effect aware using bcftools **b)** Variants called using GATK, batch effect aware and accounting for downsampling of higher coverage samples.

Figure 1 shows the geographic clustering of samples from mainland Europe and Britain, with PC2 showing greater resolution of population structure within samples from Europe. However, for the variants called without batch correction (Figure 1a) a clear effect of the sequencing run (technology) is noticeable as British and Welsh data produced on HiSeq and NovaSeq (run1) cluster together and away from the data produced through the second NovaSeq run. In contrast, Figure 1b demonstrates a greater resolution in both Welsh and English samples from different runs clustering together.

The addition of including domestic ferret samples (Figure 2) further resolves the population structure within British samples.

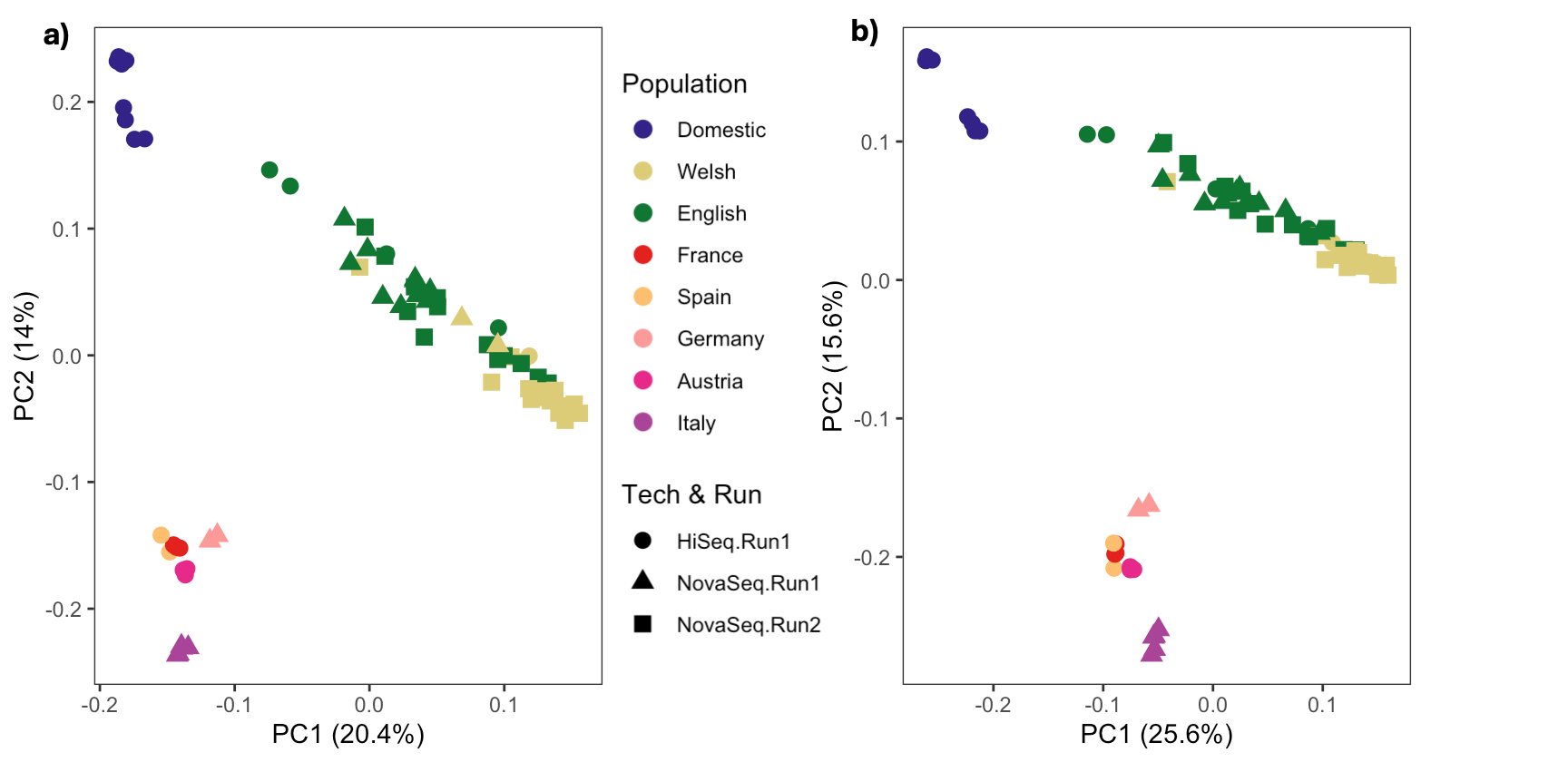

Figure 2: Principle component analysis (PCA) of whole genome SNP’s for British, Domestic ferret and European samples. Figure **a)** demonstrates the PC’s ran with SNP’s that were called with being batch effect aware using GATK, but no downsampling to account for higher coverage samples **b)** demonstrate the PC’s ran with the SNP dataset that were called using GATK, batch effect aware and accounting for downsampling of higher coverage samples.

Removing the European samples and rerunning the PCA with only British SNP data clearly shows clustering based on sequencing runs (Figure 3a). This has then been resolved using the methods above in Figure 3b.

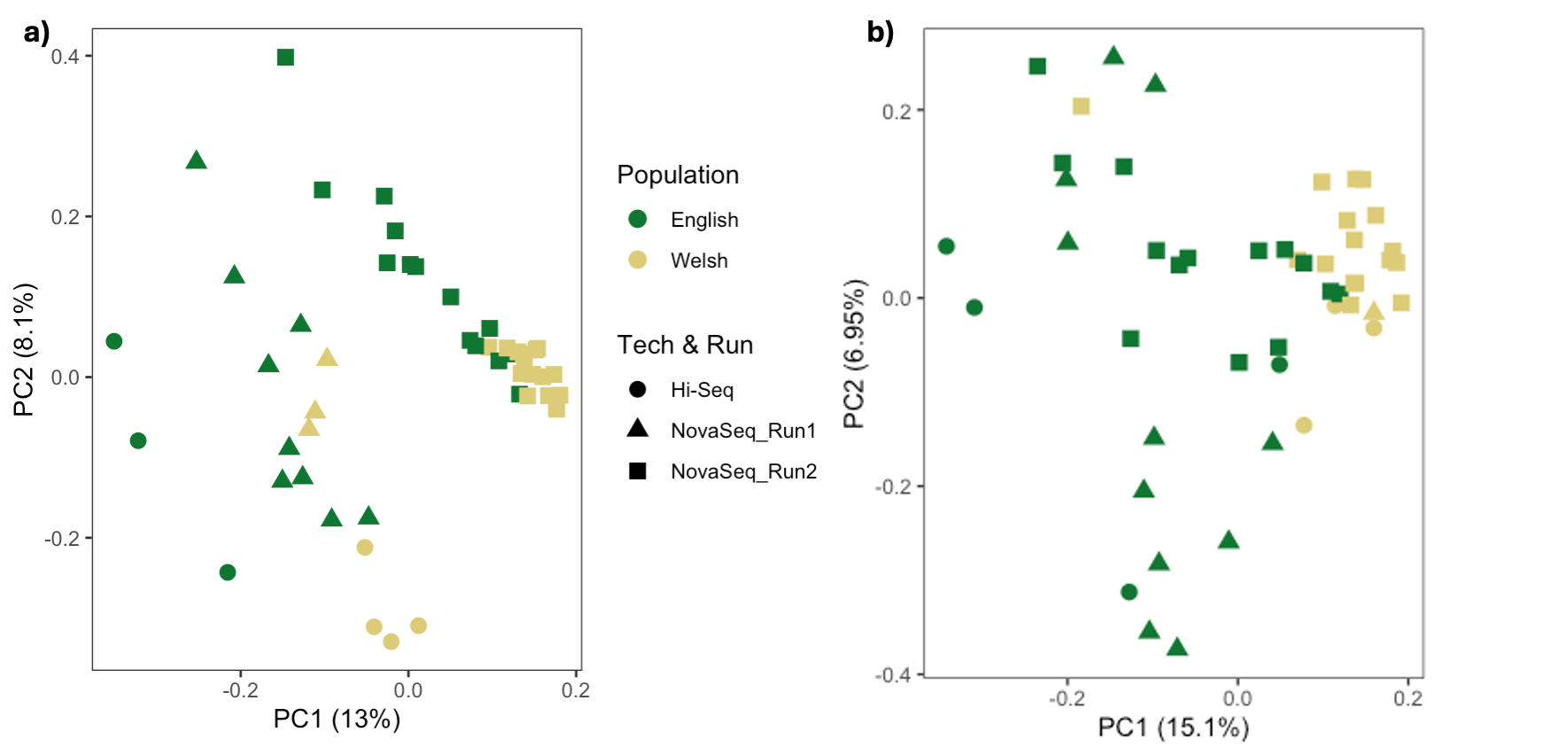

Figure 3: Principle component analysis (PCA) of whole genome SNP’s for British samples. Figure **a)** demonstrates the PC’s ran with SNP’s that were called without being batch effect aware using bcftools **b)** demonstrate the PC’s ran with the SNP dataset that were called using GATK, batch effect aware and accounting for downsampling of high coverage samples.

**References**

Auwera, Geraldine van der, and Brian D. O’Connor. 2020. *Genomics in the Cloud: Using Docker, GATK, and WDL in Terra*. Sebastopol, CA: O’Reilly Media.

Bolger, Anthony M., Marc Lohse, and Bjoern Usadel. 2014. “Trimmomatic: A Flexible Trimmer for Illumina Sequence Data.” *Bioinformatics (Oxford, England)* 30 (15): 2114–20.

Leek, Jeffrey T., Robert B. Scharpf, Héctor Corrada Bravo, David Simcha, Benjamin Langmead, W. Evan Johnson, Donald Geman, Keith Baggerly, and Rafael A. Irizarry. 2010. “Tackling the Widespread and Critical Impact of Batch Effects in High-Throughput Data.” *Nature Reviews. Genetics* 11 (10): 733–39.

Leigh, D. M., H. E. L. Lischer, C. Grossen, and L. F. Keller. 2018. “Batch Effects in a Multiyear Sequencing Study: False Biological Trends due to Changes in Read Lengths.” *Molecular Ecology Resources* 18 (4): 778–88.

Li, Heng, and Richard Durbin. 2009. “Fast and Accurate Short Read Alignment with Burrows-Wheeler Transform.” *Bioinformatics (Oxford, England)* 25 (14): 1754–60.

Lou, Runyang Nicolas, and Nina Overgaard Therkildsen. 2022. “Batch Effects in Population Genomic Studies with Low-Coverage Whole Genome Sequencing Data: Causes, Detection and Mitigation.” *Molecular Ecology Resources* 22 (5): 1678–92.

McKenna, Aaron, Matthew Hanna, Eric Banks, Andrey Sivachenko, Kristian Cibulskis, Andrew Kernytsky, Kiran Garimella, et al. 2010. “The Genome Analysis Toolkit: A MapReduce Framework for Analyzing next-Generation DNA Sequencing Data.” *Genome Research* 20 (9): 1297–1303.
